# Iterative Subtractive Binning of Freshwater Chronoseries Metagenomes Identifies of over Four Hundred Novel Species and their Ecologic Preferences

**DOI:** 10.1101/826941

**Authors:** LM Rodriguez-R, D Tsementzi, C Luo, KT Konstantinidis

## Abstract

Recent advances in sequencing technology and accompanying bioinformatic pipelines have allowed unprecedented access to the genomes of yet-uncultivated microorganisms from a wide array of natural and engineered environments. However, the catalogue of available genomes from uncultivated freshwater microbial populations remains limited, and most genome recovery attempts in freshwater ecosystems have only targeted few specific taxa. Here, we present a novel genome recovery pipeline, which incorporates iterative subtractive binning and apply it to a time series of metagenomic datasets from seven connected locations along the Chattahoochee River (Southeastern USA). Our set of Metagenome-Assembled Genomes (MAGs) represents over four hundred genomospecies yet to be named, which substantially increase the number of high-quality MAGs from freshwater lakes and represent about half of the total microbial community sampled. We propose names for two novel species that were represented by high-quality MAGs: “*Candidatus* Elulimicrobium humile” (“*Ca*. Elulimicrobiota” in the “Patescibacteria” group) and “*Candidatus* Aquidulcis frankliniae” (“Chloroflexi”). To evaluate the prevalence of these species in the chronoseries, we introduce novel approaches to estimate relative abundance and a habitat-preference score that control for uneven quality of the genomes and sample representation. Using these metrics, we demonstrate a high degree of habitat-specialization and endemicity for most genomospecies observed in the Chattahoochee lacustrine ecosystem, as well as wider species ecological ranges associated with smaller genomes and higher coding densities, indicating an overall advantage of smaller, more compact genomes for cosmopolitan distributions.

## Introduction

Freshwater environments represent a major microbial habitat on Earth, hosting an estimated 1.3×10^26^ prokaryotic cells worldwide [1, 2]. The level of diversity in microbial freshwater communities is orders of magnitude lower than that of other major environments such as soil and seawater [3], making them a tractable but globally important model for studying microbial community ecology. However, the lack of comprehensive sets of reference genomes and low cultivation rates hinder the study of these communities. On average, a quarter of freshwater community members detected by 16S rRNA gene or metagenomic surveys belong to yet-uncultured phyla, with an additional two thirds belonging to uncultured genera, families, or classes [4]. In fact, only a tenth of freshwater microbial cells belong to cultivated species or genera, the smallest cultivated fraction among all major environments on Earth (*i.e.*, environments with over 10^25^ microbial cells estimated worldwide [4]; but see also [5]). Recent efforts to recover metagenome-assembled genomes (MAGs) from freshwater environments have largely targeted specific taxa [6–10]. A few recent attempts recovered MAGs from all *Bacteria* and *Archaea* present in freshwater communities and resulted in three collections of MAGs from a lake in Siberia (Lake Baikal) and three lakes in North America (Lake Mendota, Trout Bog Lake, and Upper Mystic Lake) [11–13], as well as two collections from rivers in India (Ganges River) and Greece (Kalamas River) [14, 15]. The fraction of the communities captured by these MAGs or other reference genomes is typically moderate to low due to the high diversity of freshwater communities as well as the limitations of the underlying binning methods, which are not optimized for chronoseries datasets from natural habitats but rather for single or small sets of samples from the exact same microbial community. Temporal and spatial series from freshwater ecosystems are even sparser; yet, such data could provide a more complete picture of seasonal and biogeographic patterns of the corresponding microbial communities that are important for human activities.

We introduce here a pipeline for the recovery of MAGs from sets of metagenomes through iterative subtractive binning and apply it to a metagenomic chronoseries from freshwater lakes and estuaries along the Chattahoochee River (Southeast USA). The abundance distribution of these population genomes in the meta-community was studied using two methodological innovations: an estimation of relative abundance controlling for completeness and micro-diversity issues in the genomes, and an ecologic preference score controlling for uneven sample representation. The collection of MAGs presented here captures 50-60% of the total source communities, which is about three times larger than previous binning efforts from comparable freshwater environments, and includes representatives from taxa yet to be named, ranging from novel species of previously described genera to novel phyla.

## Materials and Methods

Additional information on software versions and parameters used is available in Table 1, and additional details are provided in the Text S1.

**Table 1:**
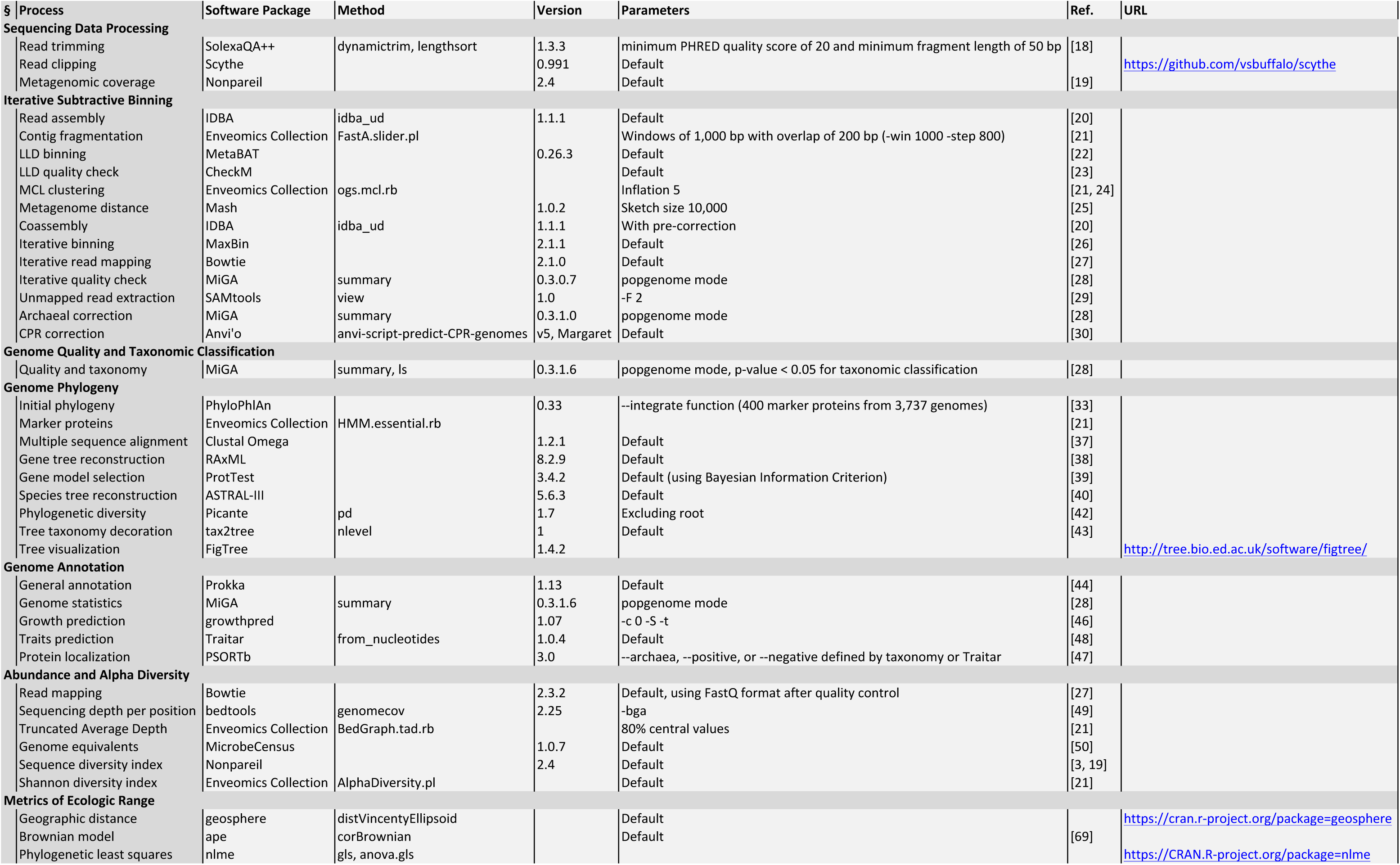
Software used in this study, sorted by method sections.

### Sample Collection and Metagenomic Sequencing

All samples were collected from the lower epilimnion (typically 3-5m depth) of the Southeastern U.S. Lakes Lanier (GA), West Point (GA/AL), Harding (GA/AL), Eufaula (GA/AL), and Seminole (GA/FL) at least 10 m away from the littoral zone, and two locations in the Apalachicola estuary, off the coasts of Apalachicola and East Point (FL). Water samples were immediately stored at 4°C and processed typically within 1-4 h, and no more than a day post collection. Water was sequentially filtered with a peristaltic pump through 2.5 µm and 1.6 µm porosity glass microfiber filters (Whatman), to capture large particles and eukaryotic cells, and microbial cells were eventually retained on 0.2 µm porosity Sterivex filters (Millipore). Thus, all sequenced metagenomes represent the 1.6-0.2 µm cell size fraction, except LLGFA_1308A and LLGFA_1309A that represent the 2.5-1.6 µm fraction. Filters were preserved at -80°C. DNA extraction was performed as previously described [16] with minor modifications and samples were sequenced using Illumina MiSeq and HiSeq sequencers (see Text S1, Metagenomic Sequencing). In addition, we included in our metagenome collection previously obtained viral enrichments (viral metagenomes) from the same freshwater samples [17] that were found to be highly contaminated with bacterial cells. Those viral metagenomes were included in the binning process, but not in subsequent analyses.

### Sequencing Data Processing

All sequenced metagenomic datasets were subjected to quality control and those not passing minimum requirements were re-sequenced. Sequencing reads were trimmed and clipped using SolexaQA++ [18] and Scythe. Abundance-weighted average coverage of the datasets was estimated using Nonpareil [19]. A minimum dataset size of 1Gbp after trimming and 50% coverage were required for all samples in this study (Table S1).

### Iterative Subtractive Binning

An initial binning methodology was implemented using metadata-dependent grouping of samples to recover high-quality metagenome-assembled genomes (MAGs; Fig. 1, top row). Specifically, we grouped and co-assembled all cell-metagenomic samples from Lake Lanier (34 samples, 120 Gbp in total). The co-assembly strategy consisted of initial individual assemblies (IDBA-UD [20]), cutting resulting contigs (FastA.slider.pl [21]), and reassembling the fragments from all samples (IDBA-UD). We binned the final contigs using MetaBAT [22] and evaluated genome quality with CheckM [23]. MAGs with estimated completeness above 75% and contamination below 5% were considered of high quality, and the resulting set was labeled **LLD**.

**Figure 1:**
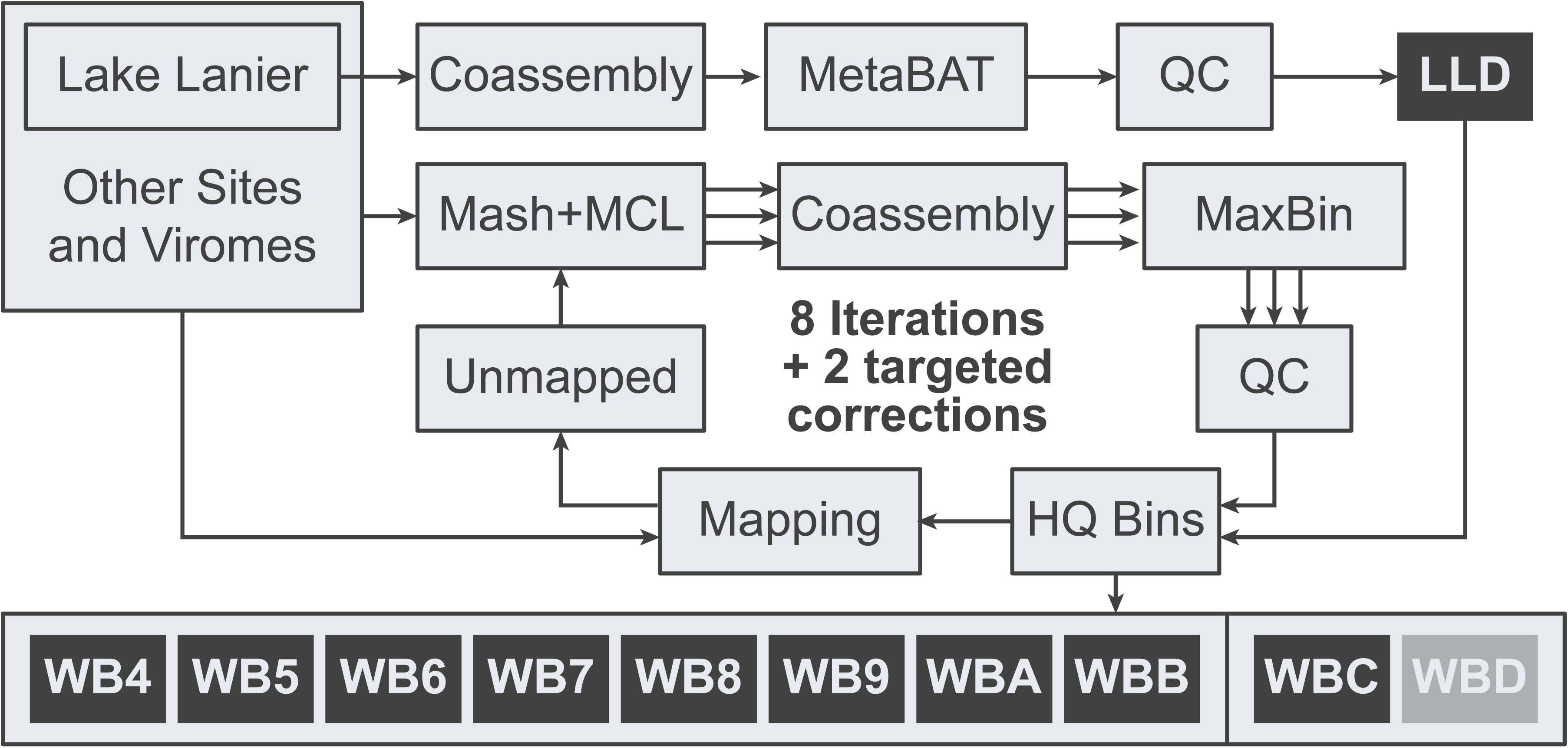
Diagram of the iterative subtractive binning methodology applied in this study. Input data (bold) and processes are depicted as light grey boxes, data flow as arrows, and output sets of MAGs as dark grey boxes. The initial non-iterative binning of Lake Lanier metagenomes corresponds to the set LLD, and the 8 iterations including all datasets correspond to the sets WB4-WBB. After the iterative approach, two targeted corrections were applied corresponding to WBC (*Archaea*) and the empty set WBD (CPR). QC stands for Quality Control, and HQ stands for High Quality.

Next, we implemented a strategy to recover MAGs using the complete collection of samples (Fig. 1). Our samples consisted of a roughly continuous two-dimensional scheme (temporal/spatial components), making metadata-based grouping of samples prone to subjective calls. Instead, we performed a sequence-based grouping by Markov Clustering (MCL) [21, 24] of Mash distances [25] using only values below 0.1. Each group was co-assembled (IDBA-UD), binned (MaxBin [26], Bowtie [27]), and evaluated using MiGA [28]. MAGs with estimated genome quality above 50 were considered of high quality (see below genome quality definition), and the first resulting set was labeled **WB4**. The resulting set of high-quality MAGs (LLD + WB4) was used as reference database to map reads from all samples (Bowtie), and unmapped reads (SAMtools [29]) were used as input for Mash/MCL clustering, iterating the process described above to produce sets **WB5-WBB** (Fig. 1). The number of iterations was determined by saturation of phylogenetic breadth and fraction of reads mapping (Fig. 2). Finally, two corrections were implemented targeting groups that typically generate quality underestimations. First, a correction for archaeal genomes in MiGA was used to recover high-quality genomes from *Archaea* in all iterations (**WBC**). Second, the Random-Forest classifier for Candidate Phyla Radiation (CPR) scripts in Anvi’o [30] were used to detect high-quality genomes from CPR in all iterations, which didn’t yield any additional MAGs. The complete collection of high-quality MAGs is hereafter designated WB (Table S2).

**Figure 2:**
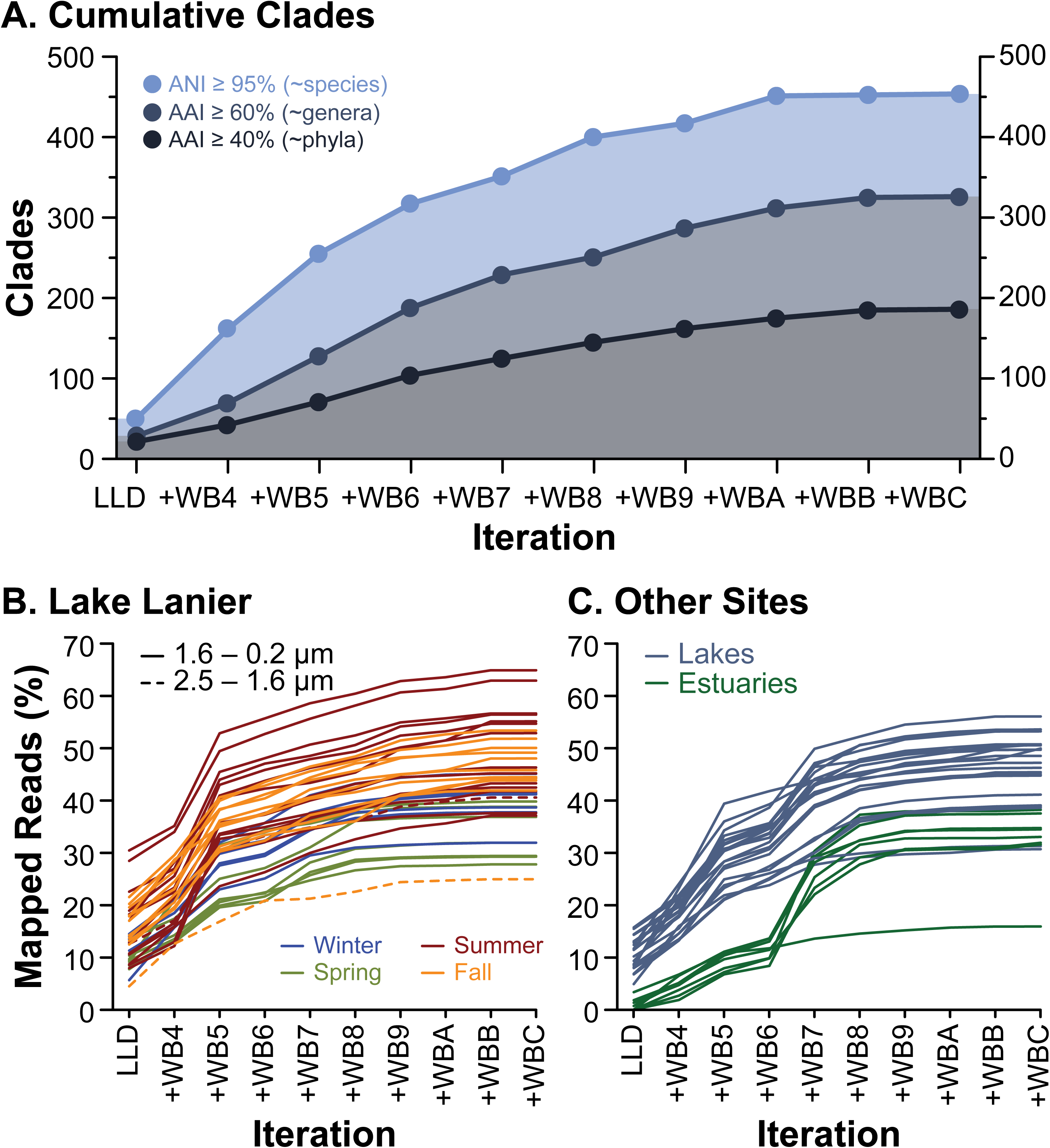
Saturation of captured diversity along the iterative subtractive binning rounds. **(A)** Total number of clades captured with ANI ≥ 95% (light blue, representing species level), AAI ≥ 60% (dark blue, roughly corresponding to genus level), and AAI ≥ 40% (grey, roughly corresponding to phylum level). Note that the range of AAI values (a proxy for genetic relatedness) within genera and phyla typically varies between clades, and the latter two thresholds shouldn’t be considered as precise estimates of taxonomic diversity. **(B-C)** Total fraction of metagenomic reads from each dataset mapping to the complete (cumulative) collection of MAGs after each iteration. Each line represents a metagenomic dataset derived from Lake Lanier (B), other lakes (C, blue), or estuarine samples (C, green).

### Genome Quality and Taxonomic Classification

The quality and taxonomic classification of MAGs were evaluated using MiGA. Briefly, a composite index of genome quality was used, defined as “Completeness - 5×Contamination”, where both completeness and contamination were determined by the presence and copy number of genes typically found in genomes of *Archaea* and *Bacteria* in single copy [21, 28]. Taxonomy was determined by MiGA with the NCBI Genome Database, Prokaryotic section (henceforth NCBI_Prok; MiGA Online; Jan-2019) [28]. MiGA also performs a de-replication of the collection by generating groups of genomes with ANI ≥ 95% using ogs.mcl.rb [21, 24]. These clusters, analogous to bacterial or archaeal species [31, 32] are hereafter termed genomospecies (gspp, singular gsp).

### Genome Phylogeny

Two phylogenetic approaches were used to place the obtained MAGs in the context of the tree of *Bacteria* (only 4 distinct species of *Archaea* were recovered). First, we used PhyloPhlAn [33] to place the genomes in the context of a general-purpose widely used genome collection. Next, we generated a phylogenetic reconstruction using the high-quality MAGs in this study classified as *Bacteria*, and all best-match entries (highest AAI) of our set against five collections of genomes available in MiGA Online at http://microbial-genomes.org/projects. Namely, a manually curated collection of MAGs from various projects (**GCE**), a set of MAGs recovered from the Tara Oceans expedition (**TARA**) [34], a collection of MAGs recovered from various environments excluding human microbiome (**UBA**) [35], all complete genomes available in NCBI (**NCBI_Prok**), and all available genomes (complete or draft) from type material (**TypeMat**) [36]. Marker proteins were extracted from all the abovementioned genomes (HMM.essential.rb [21]), and those present in at least 80% of the genomes were selected and independently aligned (Clustal Omega [37]). Next, maximum likelihood gene trees were constructed for individual alignments using RAxML [38] with model selected using ProtTest [39]. Finally, a species tree was estimated from the best-scoring ML trees reconstructed for each gene using ASTRAL-III [40].

Both final trees (PhyloPhlAn and ASTRAL) were used to estimate phylogenetic gain for the WB collection using Faith’s Phylogenetic Diversity (PD [41]; Picante [42]):

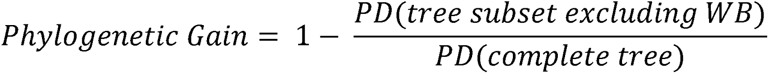

In the ASTRAL tree, branch lengths for all terminal nodes were set to zero in this analysis. The taxonomic classification reported by NCBI for the genomes in the collections TypeMat, NCBI_Prok, and UBA was recovered by MiGA, and used to calibrate taxonomic limits in coalescent units by identifying the median values between taxonomic ranks. In addition, this taxonomic information was used to decorate the rooted ASTRAL species tree (tax2tree [43]). The tree was visualized using FigTree.

### Genome annotation

Functional annotation of all genomes was performed using Prokka [44]. Protein annotations from COG (Cluster of Orthologous Groups of proteins) were mapped to COG categories using eggNOG [45]. Gene coding density, G+C content, and other descriptive statistics, as well as genome completeness, contamination, and quality were calculated using MiGA [28]. Growth rate and optimal growth temperature were predicted using growthpred [46]. Extracellular proteins were predicted using PSORTb with Gram staining predicted by Traitar [47, 48].

### Abundance and Alpha Diversity

The abundance of each gsp was estimated using the MAG of highest genome quality as representative. For each metagenomic dataset, the sequencing depth was estimated per position (Bowtie [27], bedtools [49]) and truncated to the central 80% (BedGraph.tad.rb [21]), a metric hereafter termed **TAD** (truncated <u>a</u>verage sequencing <u>d</u>epth). Abundance was estimated as TAD normalized by the genome equivalents of the metagenomic dataset (MicrobeCensus [50]), resulting in units of community fraction. A gsp was considered to be present in a sample if the TAD was non-zero (equivalent to sequencing breadth ≥ 10%, previously shown to correspond to confidence of presence > 95% [51]). The alpha-diversity was estimated using the sequence diversity *N_d_* projected to Shannon diversity *H’* (Nonpareil [3]), as well as *H’* on the gspp abundance profile (AlphaDiversity.pl [21]). Additional details are available on Text S2.

### Preference Scores

In order to determine the preferential presence of a gsp in a given set of samples while accounting for the geographic and environmental biases in the dataset collection used here, we devised a preference score accounting for the expected abundance of a gsp in a given dataset (see Text S1, Preference Score). Briefly, we first estimated the **observed bias** (*i.e.*, over- or under-representation) in presence frequency of a gsp in a given set of samples compared to the rest of the samples. Next, we estimated the **expected bias** assuming that there is no preference by normalizing by both gsp presence frequency across all samples as well as the presence frequency of all gspp in each sample. This is achieved by estimating the expected frequency of each MAG in a metagenome as the frequency of MAGs in that metagenome multiplied by the frequency with which the MAG is observed across metagenomes. Finally, we calculate the ratio of these two biases (observed/expected) maintaining the sign of the observed bias. The preference score of gsp *i* for sample set *t* is termed 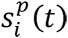. A score was considered significant when 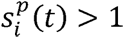 (preference for the set *t*) or 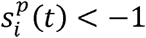 (preference against set *t*). No clear preference was established for gspp with 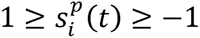.

### Samples from Other Projects

In addition to the metagenomes sequenced as part of our study, we used previously reported metagenomes from other sites and environments for comparisons. These metagenomes, derived from previous studies [13, 52–65], or recovered via MGnify [66], are described in Table S3. The raw reads were obtained from the European Nucleotide Archive (EBI ENA) and processed as described above. The metadata for each sample was obtained from EBI ENA or the original studies, including biome, aquatic habitat, and geographic location (latitude and longitude).

### Metrics of Ecologic Range

Ecologic ranges were measured in different dimensions reflecting environment and geographic location. Environments were characterized by **biome** (one of brackish water, estuary, estuary sediment, freshwater sediment, glacier, groundwater, human gut, lake, marine oxygen minimum zone, marine surface, marine water column, river, or soil) or **aquatic habitats** (brackish, estuary, freshwater, marine, non-aquatic), and for each gsp the **count breadth** (number of biomes or aquatic habitats) was determined by presence as non-zero TAD in the corresponding samples of the biome or habitat. In addition, the frequency of presence of a gsp across samples per biome or aquatic habitat was used to estimate the entropy (natural units), as proposed by Levins [67] (**unweighted Levins’ breadth**). In order to account for the estimated abundances (and not only inferred presence), we also defined average abundance across samples per biome or aquatic habitat to estimate entropy (**weighted Levins’ breadth**). Geographic distances were estimated using the distance on the ellipsoid [68] (geosphere). For each gsp, two geographic ranges were estimated: the maximum distance between any two samples where the gsp is present (**geodesic range**), and the maximum latitudinal range of samples where the gsp is present (**latitude range**). Correlations between traits and ecologic ranges were evaluated by Pearson’s linear correlation for continuous variables and Spearman’s rank correlation for counts. Additionally, correlations along the ASTRAL phylogenetic reconstruction were evaluated using phylogenetic generalized least squares (nlme) assuming a Brownian model (ape [69]).

## Results

### Freshwater Metagenomic Datasets

We sequenced a total of 69 metagenomic datasets derived from water samples from Lakes Lanier, Harding, Eufaula, and Seminole, and the estuarine locations of Apalachicola and East Point along the Chattahoochee River, in the Southeastern continental USA (Table S1). All samples were collected from the lower epilimnion to allow comparisons across sites. All samples were required to have at least 60% coverage as estimated by Nonpareil [3], except for LL_1007C (46% coverage) that had a high-coverage replicate (LL_1007B, 83% coverage). Excluding the latter (LL_1007C), samples had an average community coverage of 76% (Inter-Quartile Range –IQR–: 70.6-81.3%) and an average total size after trimming of 3.4 Gbp (IQR: 2.6-4.4 Gbp). The sequence diversity estimated by Nonpareil (*N_d_*) was on average 19.6 (IQR: 19.3-20.0), typical of freshwater microbial communities [3].

### Iterative Subtractive Binning

An iterative subtractive binning methodology was applied to the collection of metagenomes described here. Briefly, metagenomic datasets were processed by grouping metagenomes by read-level similarity (Mash distances), co-assembling with or without subsampling, binning, mapping reads to high-quality obtained MAGs, and iterating this methodology with the resulting unmapped sequencing reads (Fig. 1; see also Materials and Methods). This method produced a total of 1,126 MAGs grouped in 462 genomospecies, *i.e.*, clusters with intra-cluster ANI ≥ 95%. The average estimated completeness of the MAGs in this set was 75.4% (IQR: 66.7-84.7%), and the average estimated contamination was 2.10% (IQR: 0.9-2.7%). This result contrasts with the 199 MAGs identified in the initial non-iterative binning (LLD), grouped in 166 gspp (Fig. 2-A), indicating that the iteration process captured at least three times higher taxonomic diversity. The initial quality control excluded all archaeal genomes captured, and the archaeal correction (WBC) recovered 22 genomes from 4 gspp. No additional genomes were recovered by the CPR correction.

### Diversity Captured

The initial non-iterative binning (LLD) captured only 8-14% (IQR; average: 11.5%) of the total metagenomic reads, depending on the dataset considered, whereas the final set captured 38-50% (IQR; average: 43.2%) of the total metagenomic reads (Fig. 2-B-C). These figures underscore the large increase in representation of the community throughout the iterative process. However, it is expected that this representation be strongly biased towards the most abundant members of the community. In order to reduce the effects of genome size variation, completeness, and other artifacts, we estimated relative abundance of MAGs as truncated sequencing depth (TAD) normalized by genome equivalents (see Methods and Text S2). The estimated fraction of the community captured by the final set of MAGs was 42-59% (IQR; average: 50.2%). Importantly, this fraction is considerably larger than that of other available MAG sets from freshwater lakes, further underscoring the usefulness of iterative subtractive binning. For instance, a previous study on the microbial communities of Upper Mystic Lake (Massachusetts, USA) [12] recovered a set of 87 genomes from 14 metagenomic datasets. Using the same abundance estimations as above, we calculated that those 87 genomes captured 11-18% (IQR; average: 14.9%) of the source communities. A smaller set of 35 MAGs recovered from two metagenomic dataset from the waters under the surface ice layer of Lake Baikal (Siberia, Russia) [11], resulted in 10 and 9.7% of the source communities captured at 20- and 4-m-deep samples, respectively. Finally, a set of 194 MAGs recovered from three chronoseries from the eutrophic Lake Mendota and the humic Trout Bog Lake (Wisconsin, USA) [13] resulted in 20.2%, 31.9%, and 38.4% of the communities captured in Lake Mendota, and the epilimnion and hypolimnion of Trout Bog Lake, respectively. In addition, we evaluated two riverine MAG datasets from Rivers Ganges (India) and Kalamas (Greece). The former, composed of 104 MAGs, captured on average 23.6% of the source communities (IQR: 18-33%), and the latter with 14 MAGs captured 7.4% (IQR: 6-10%). Overall, freshwater MAG sets from previous studies captured on average 16.7% of the source communities (IQR: 10-20%), about three times less than the WB set presented in this study. However, note that the high metagenomic read recovery from the WB collection does not preclude other biases for community-level diversity assessment. Most notably, we identified that MAGs capture a disproportionally larger fraction of less diverse communities, indicating that profile summary statistics such as Shannon diversity or Richness estimations should not be computed directly from collections of MAGs (Fig. S1, Text S2).

### Phylogenetic Diversity and Novelty

We reconstructed a coalescent-based phylogeny of all high-quality bacterial MAGs in this study (n=1,108 in 462 gspp) and related genomes (best-hit by AAI) in different reference collections (Fig. 3). The best-hit sets included genomes from GCE (n=96, from 591 genomes/393 gspp), TARA (n=173, from 957 genomes/856 gspp), UBA (n=224, from 7,903 genomes/4,042 gspp), NCBI_Prok (n=226, from 13,826 genomes/4,271 gspp), and TypeMat (n=143, from 9,077 genomes/6,939 gspp). Marker proteins from all the abovementioned genomes (n=1,970) present in at least 80% of the genomes were selected (n=70, from 110 proteins evaluated) for gene-tree reconstructions reconciled in the final species tree.

**Figure 3:**
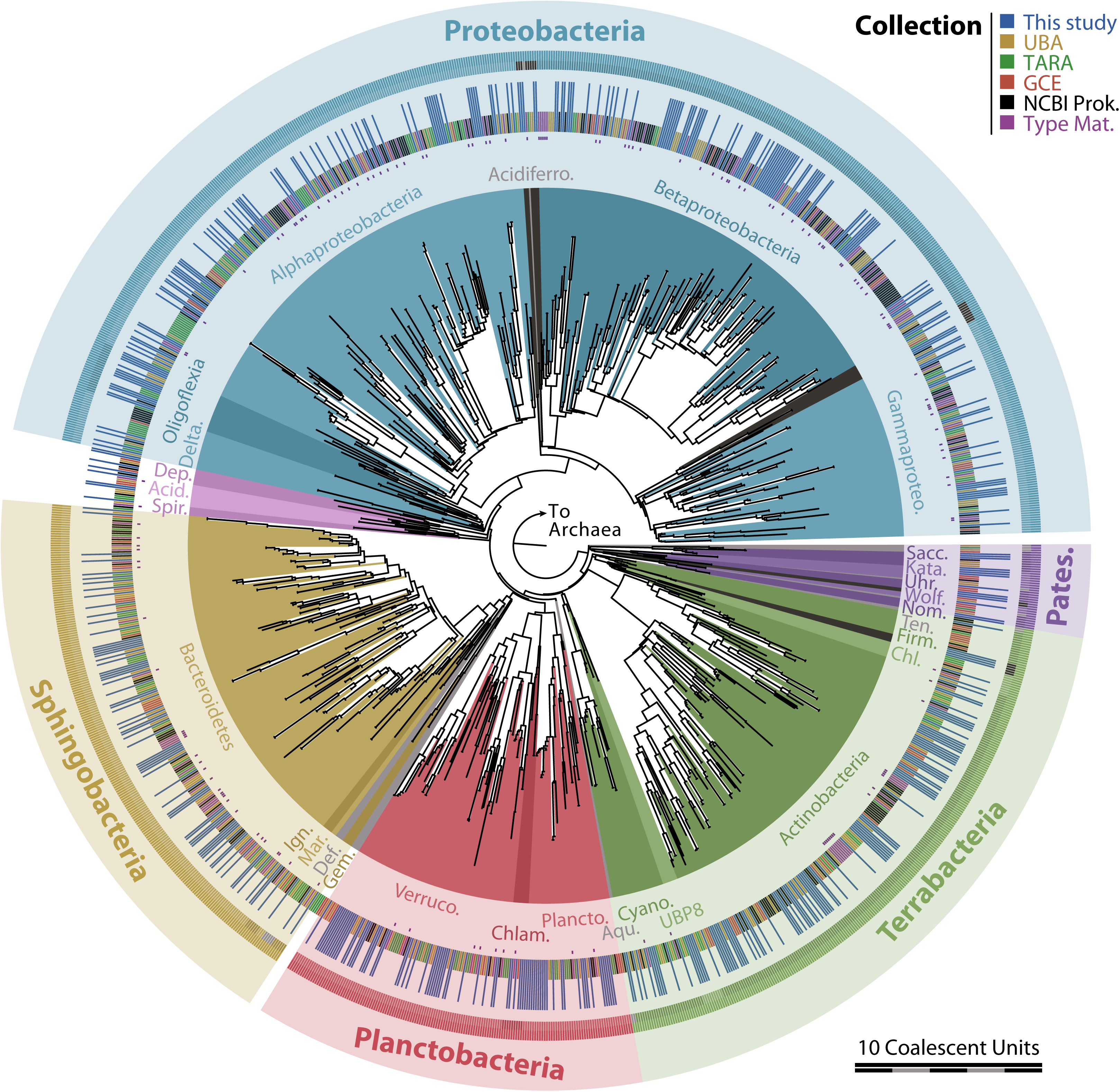
Phylogenetic reconstruction of the bacterial MAGs in this study and closest relatives derived from five different genome collections. The phylogeny was reconstructed using coalescent-based species tree estimation [40] from 70 gene trees reconstructed by Maximum Likelihood [38, 39]. The tree is decorated with colored backgrounds corresponding to phyla (or classes in “Proteobacteria”), labeled in the innermost ring. Light grey background corresponds to taxa not including any representatives from our collection, and dark grey corresponds to yet-unnamed taxa. The next ring indicates the genome collection (see legend), emphasizing genomes from type material (purple, with accent dots inwards) and from the current study (blue, extending outwards). The following double-ring corresponds to the innermost background (phyla or classes, inwards) and the larger containing group as labeled in the outermost ring (superphyla or the phylum “Proteobacteria”, outwards). The labels use abbreviations for the following taxa (clockwise): “Patescibacteria” (Patesc., also referred to as CPR), “*Ca*. Saccharibacteria” (Sacc.), “*Ca*. Katanobacteria” (Kata., also referred to as WWE3), “*Ca*. Uhrbacteria” (Uhr.), “*Ca*. Wolfebacteria” (Wolf.), “*Ca*. Nomurabacteria” (Nom.), “Tenericutes” (Ten.), “Firmicutes” (Firm.), “Chloroflexi” (Chl.), “Cyanobacteria” (Cyano.), “Aquificae” (Aqu.), “Planctomycetes” (Plancto.), “Chlamydiae” (Chlam.), “Verrucomicrobia” (Verruco.), “Gemmatimonadetes” (Gem.), “Deferribacteres” (Def.), “Marinimicrobia” (Mar.), “Ignavibacteriae” (Ign.), “Spirochaetes” (Spir.), “Acidobacteria” (Acid.), “Dependentiae” (Dep.), *Deltaproteobacteria* (Delta.), *Acidiferrobacteria* (Acidiferro.), and *Gammaproteobacteria* (Gammaproteo.).

We characterized the global gain in phylogenetic diversity represented by our collection with respect to two reference sets. First, in the set of best-matching genomes described above (ASTRAL tree), our collection represents about 409 novel species (out of 999 total species-level clades) and 70 novel genera (out of 332), based on approximated calibration of taxonomic ranks (as retrieved from NCBI) in the reconstructed phylogeny (Fig. S4-A-B). Overall, the gain in summed branch lengths (phylogenetic diversity) was estimated at 24.8%. A similar value of phylogenetic gain was obtained when comparing against a second reference set obtained directly from PhyloPhlAn (24.5%; Fig. S4-C-D). However, note that both estimates of phylogenetic gain are likely inflated since the former reference set does not include groups distant from any MAG in our collection (*i.e.*, we only used reference genomes identified as best matches to WB), and the latter does not include recently described taxa (PhyloPhlan version 0.99, last updated May/2013).

### Presence in Other Sites and Ecosystems

We evaluated the presence of the WB gspp in samples from different environments, mainly aquatic (Fig. 4). WB species were considered present in a sample if their sequencing depth was at least 10%, which corresponds to confidence of presence > 95% [51]. In order to determine environmental or geographic preference, we estimated preference scores based on the frequencies of presence in different sets of samples, normalizing by the baseline distribution of each gsp and the probability of capturing any gsp in a given sample, and implicitly accounting for sample size and community evenness among other factors (see Methods; Fig. 4-A). Gspp tended to cluster in two main groups: freshwater (77%) and seawater (18%), with a few gspp showing no clear preference between fresh- and seawater (4%; Fig. 5-A). From 20 gspp showing no clear preference, 19 were restricted to estuarine samples (classified as seawater in this test) and freshwater (Fig. 4-C), and one was observed in three marine samples at low abundances. Therefore, the lack of clear preference was likely the effect of low statistical power and/or water mixing in estuaries. Only 7 gspp were present in both freshwater and marine samples (6 *Synechococcaceae*), but all were detected in only 1 or 2 marine or freshwater samples at consistently low abundances (10^-5^-0.01%). Therefore, no evidence of gspp adapted to both freshwater and marine environments was found. Among those with clear freshwater preference, 73% were predominantly found in the Chattahoochee lakes, and 33 gspp (9%) displayed a preference for Lake Mendota (Fig. 5-B). Finally, 53% of the seawater gspp had a clear preference for estuarine over marine samples, whereas the rest were evenly divided in preference for marine samples or no clear preference (Fig. 5-D).

**Figure 4:**
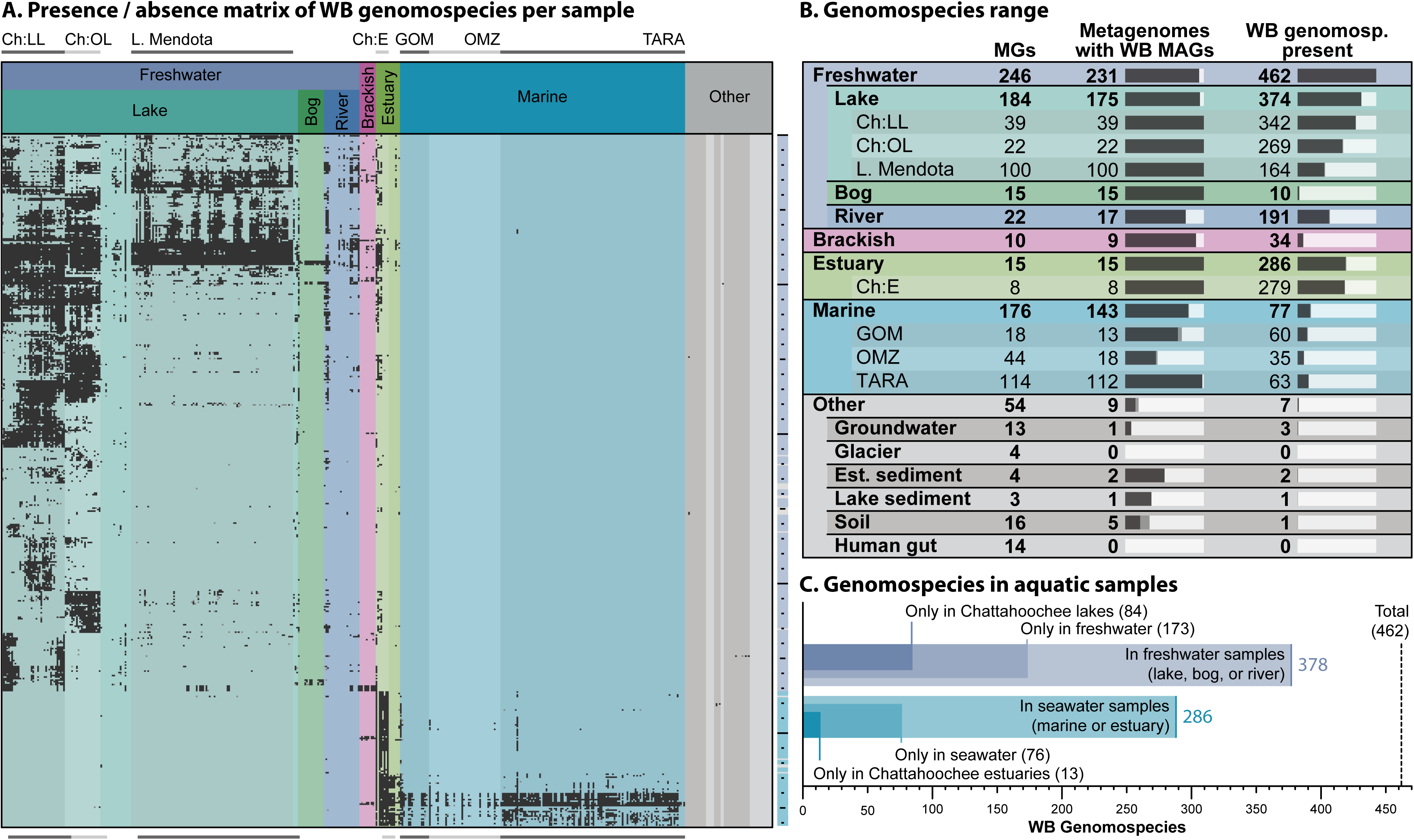
Detection of the WB genomospecies in different environments. **(A)** Presence/absence matrix of WB gspp per sample. The columns correspond to metagenomic samples, sorted by biome and source collection, and the rows correspond to WB gspp, sorted by the presence/absence pattern using Ward’s hierarchical clustering of Euclidean distances. Empty cells in the matrix correspond to TAD of zero (*i.e.*, sequencing breadth below 10%), grey cells correspond to 0 < TAD < 0.01X, and black cells correspond to TAD ≥ 0.01X. Large collections of metagenomic samples are indicated with horizontal bars at the top and bottom and matching shading in the matrix, and correspond to **Ch:LL**: Lake Lanier (Chattahoochee, this study), **Ch:OL**: other lakes from Chattahoochee (this study), **L. Mendota**: Lake Mendota (WI, USA; JGI), **Ch:E**: Estuaries from Chattahoochee (this study), **GOM:** Gulf of Mexico water column, **OMZ:** Oxygen Minimum Zone, and **TARA:** Tara Ocean expedition. For reference, the ticks on the left are spaced every 10 rows, and the marker colors correspond to gspp with freshwater preference (blue), seawater preference (teal), or no clear preference (grey; see also Fig. 5). **(B)** Summary statistics for gspp detection. Each row corresponds to a set of samples, and the columns indicate the total number of metagenomes (MGs), the number and fraction of metagenomes with WB MAGs, and the number and fraction of WB gspp present in the sample set. **(C)** Genomospecies in aquatic samples: freshwater (blue) and seawater (teal). The different marks indicate the total number of gspp in each environment (light bars) and the number of gspp found only in that environment (intermediate bars) and only in the Chattahoochee samples (dark bars).

**Figure 5:**
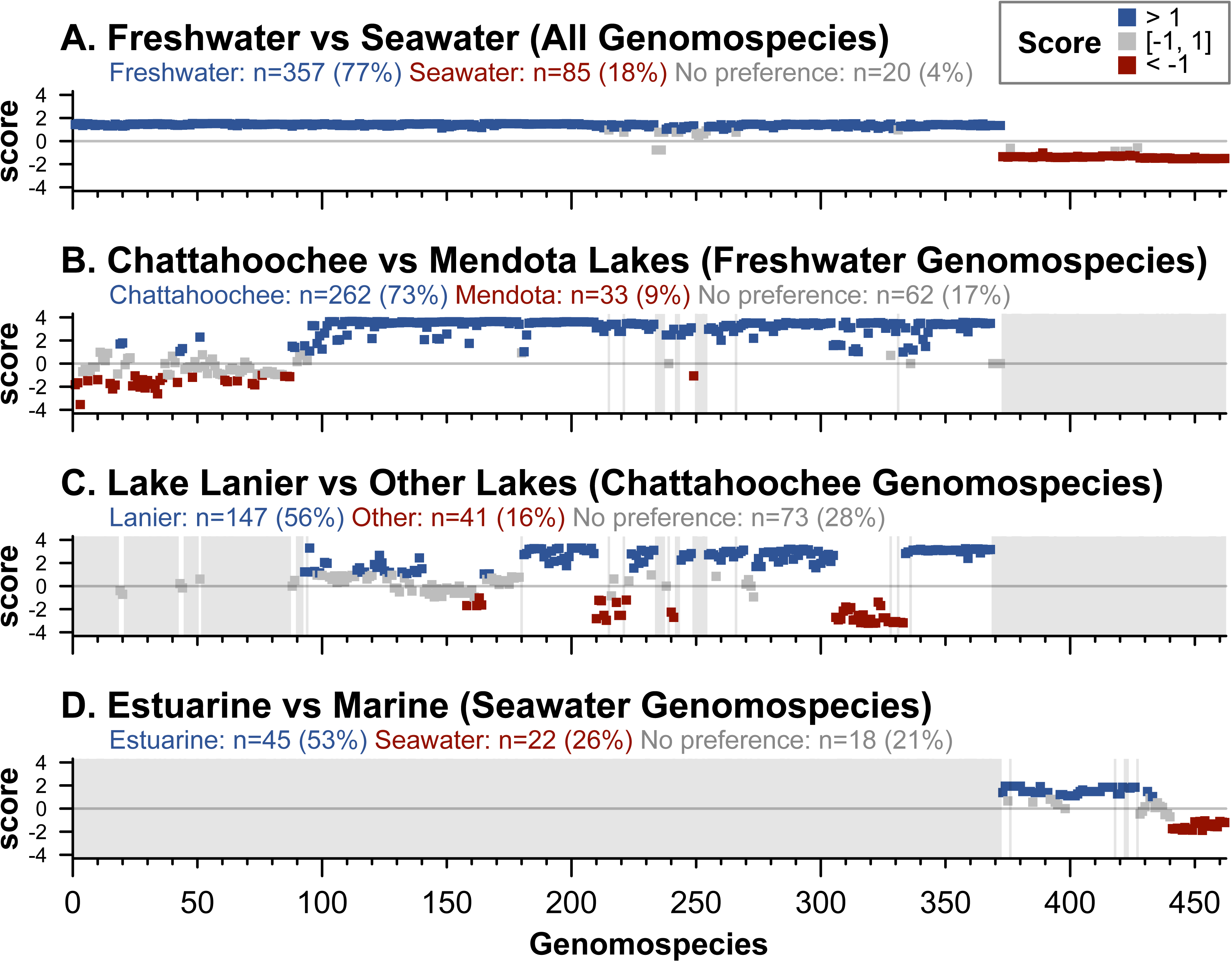
Preference scores of the WB genomospecies in different sample sets. **(A)** Preference scores for freshwater *vs.* seawater samples of all WB gspp. Larger positive values indicate stronger preference towards freshwater, and larger negative values stronger preference towards seawater. **(B)** Preference scores for Chattahoochee lakes *vs*. Lake Mendota samples among gspp with clear preference towards freshwater (blue squares in panel A). Larger positive values indicate stronger preference for Chattahoochee lakes. Shadowed areas indicate excluded gspp (without clear preference towards freshwater). **(C)** Preference scores for Lake Lanier *vs*. other Chattahoochee lakes samples among gspp with clear preference towards Chattahoochee lakes (blue squares in panel B). **(D)** Preference scores for estuarine *vs*. marine samples among gspp with clear preference towards seawater (red squares in panel A).

Next, we determined the ecologic ranges of each gsp as the number of different biomes where it could be confidently detected (**biome count**), the number of aquatic habitats (**habitat count**), the maximum geographic distance between samples where it was detected (**geodesic range**), and the maximum range of latitudes (**latitude range**). Biome and aquatic habitat breadths were additionally measured by **unweighted** (frequency of presence) and **weighted** (abundance) **Levins’ breadth** [67]. All metrics of ecologic range displayed significantly negative correlation with expected genome size (assembly length divided by estimated completeness; ρ or R between -0.18 and -0.3; p-values < 10^-5^) and positive correlation with coding density (ρ or R: 0.21-0.38; p-values < 10^-6^), indicating that more cosmopolitan and habitat-generalist gspp exhibit smaller and more compact genomes (Figs. 6, S5). Among gspp in three aquatic habitats, WB8_4xD_006 had the highest coding density (96.2%, estimated genome: 1.07Mbp), previously identified as a member of an uncharacterized clade of “*Ca.* Pelagibacterales” temporarily designated PEL8 [10]. Most gspp present in three aquatic habitats in the top 20% of coding density belong to “Actinobacteria” (n=8) or “*Ca*. Pelagibacterales” (n=4) Despite this strong taxonomic bias, correlations between coding density and ecologic ranges remained statistically significant after excluding all members of “Actinobacteria” (p-values < 10^-4^), “*Ca*. Pelagibacterales” (p-values < 2.8×10^-4^), or both (p-values < 1.4×10^-3^). On the other end, among gspp restricted to a single aquatic habitat, two genomes were particularly notable for their low coding density: WB6_1B_304 (83.49%, estimated genome: 3.39 Mbp; “Cyanobacteria”) and WB9_2_319 (85.1%, 4.77 Mbp; “Proteobacteria”; Fig. 6), and no taxonomic bias was observed in this set. In addition, genomes from more cosmopolitan gspp exhibited larger fractions of COG-annotated genes (ρ or R: 0.22-0.27; p-values < 10^-6^). This effect was possibly due to a higher prevalence of better-characterized functions (housekeeping genes, central metabolism) in smaller genomes and/or database bias towards more broadly distributed microbes. We observed a significant negative correlation of G+C% content with count breadth of aquatic habitats (R: -0.1; p-value: 0.025) and weighted Levins’ breadths of both biome and aquatic habitat (R: -0.28, -0.29; p-values: 2.5×10^-10^, 1.5×10^-9^), but not with other environmental range metrics (Fig. S5). Other genomic signatures associated with the growth strategy such as the density of ribosomal proteins (COG category J), estimated minimum generation time, and estimated optimal growth temperature were not significantly correlated with ecologic range metric (|ρ or R| < 0.07; p-values > 0.15). However, when controlling for phylogenetic relatedness (assuming correlation under a Brownian model), the minimum generation time was negatively correlated with all metrics of ecologic range (p-values < 0.035), indicating that faster growth is a trait that facilitates broader ecologic ranges among close relatives. Finally, we evaluated the possibility of larger fractions of extracellular proteins present in more cosmopolitan organisms, previously proposed as a mechanism of ecological success for pathogenic bacteria [70]. Interestingly, we observed the opposite trend: more cosmopolitan gspp were predicted to have fewer extracellular proteins as a fraction of thei genome (ρ or R < -0.17, p-values < 2.5×10^-4^).

**Figure 6:**
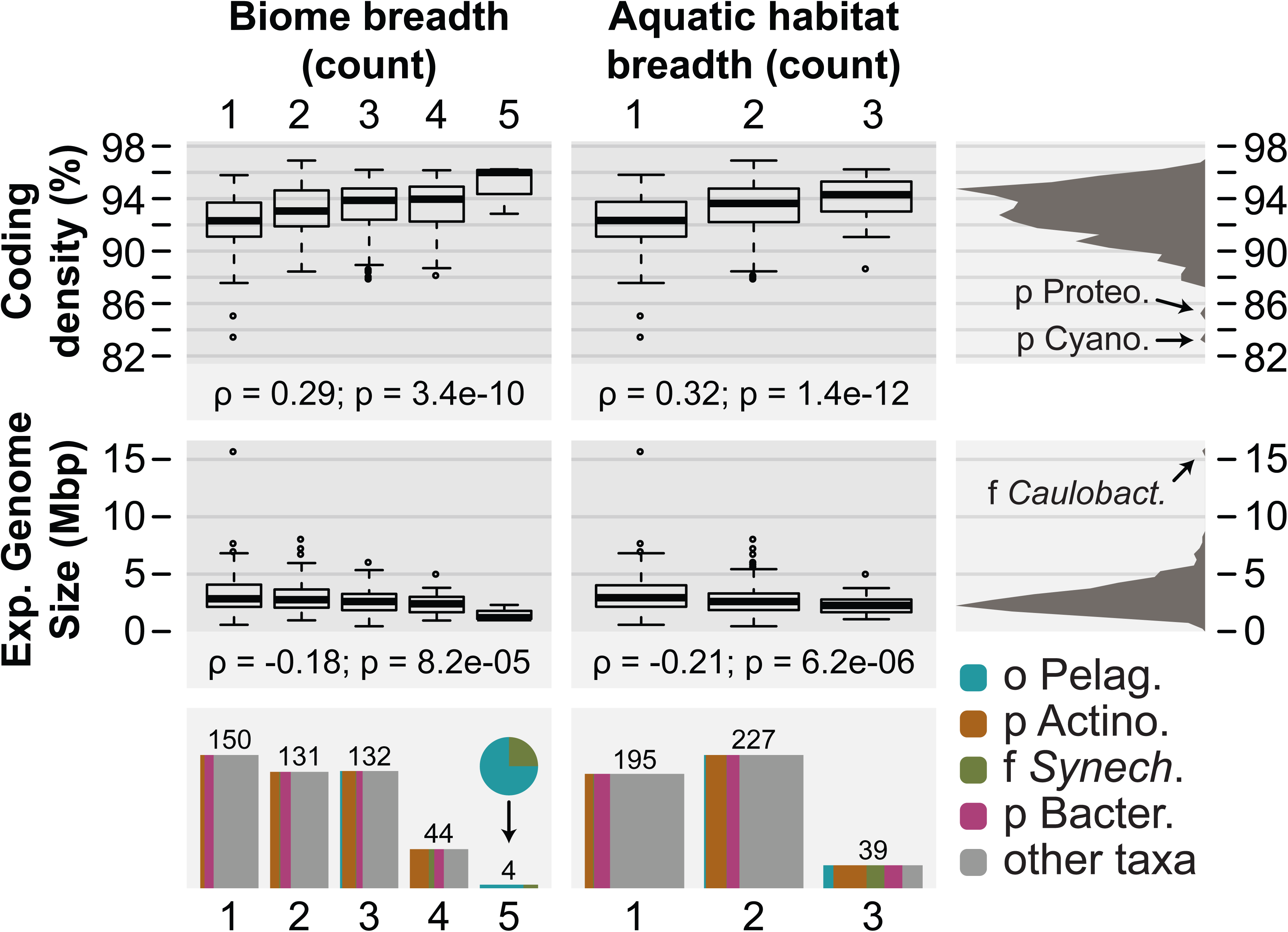
Biome and aquatic habitat breadths as functions of genome coding density and estimated size. The panels in the **top** display the coding density of the genomes for each given biome breadth (left), aquatic habitat breadth (center), and the histogram for all representative genomes (right) indicating two outliers classified in the phyla “Proteobacteria” (p Proteo.) and “Cyanobacteria” (p Cyano.) further discussed in the main text. The panels in the **middle** follow the same layout, with the rightmost histogram highlighting an outlier classified in the family *Caulobacteraceae* (f *Caulobact*.). Finally, the panels in the **bottom** indicate the distribution of genomes by biome (left) and aquatic habitat (right) breadths as bar plots with the total counts shown above each bar. Additionally, the bottom panels highlight the frequency of selected taxa that were overrepresented among cosmopolitans by the width of the colored sections (see legend), including the order “*Ca*. Pelagibacterales” (o Pelag.), the phylum “Actinobacteria” (p Actino.), the family *Synechococcaceae* (f *Synech*.), and the phylum “Bacteroidetes” (p Bacter.).

### Description of Novel Taxa

Finally, we characterized the genomes representing two novel taxa. We propose the names “*Candidatus* Elulimicrobium humile” gen. nov. sp. nov., represented by WB6_2A_207 (GenBank: RGCK00000000), from a novel phylum (“*Candidatus* Elulota” phy. nov.) within the “Patescibacteria” group, and “*Candidatus* Aquadulcis frankliniae” gen. nov. sp. nov., represented by WB4_1_0576 (GenBank: RFPZ00000000), from a novel genus within the recently described class “*Candidatus* Limnocylindria” [6] (“Chloroflexi”). Additional description of these taxa including protologues is available as Supplementary Material (Text S3 and Fig. S2).

## Discussion

In this study, we introduced a methodology for iterative subtractive binning of metagenomic collections including *de novo* grouping of samples (*i.e.*, independent of metadata) and the gradual reduction of dataset diversity for the recovery of genomes from populations with vastly different relative abundances (Fig. 1). The genomes recovered showed on average a maximum relative abundance across samples of only 0.59% of the total microbial community (IQR: 0.12-0.55%), with as many as 17% of the recovered genomospecies consistently below 0.1% relative abundance, considered the rare fraction in this ecosystem [71]. We were able to reconstruct the genome of a “Patescibacteria” bacterium for which we propose the name “*Ca*. Elulimicrobium humile”, representing a novel phylum (“*Ca*. Elulota”), that appears to be regionally widespread and endemic, but had consistently low abundance in our metagenome series (≤ 0.12%). Combined, all the gspp in our collection represent about 50% of the entire communities (Chattahoochee metagenomes), about three times more than other binning efforts in freshwater habitats. Importantly, we demonstrate that MAGs capture a larger fraction of less diverse communities. Therefore, we recommend against using summaries of abundance profiles from MAGs to characterize and/or compare entire communities (*e.g.*, measuring richness or alpha/beta diversity from MAG profiles), and emphasize the advantages on phylogenetic novelty of individual populations instead.

Overall, from 462 genomospecies detected here, 452 (98%) represent novel species on the basis of ANI or 409 (88%) on the basis of approximate phylogenetic calibration, indicating that the great majority of genomes recovered here are novel. In terms of phylogenetic novelty, about one fourth of the branch lengths of a phylogenetic reconstruction including all best matches from complete genomes, type material, and MAGs, were uniquely derived from our set (Fig. S4). Moreover, the species detected in our samples span a variety of geographic ranges, from highly restricted locally to regionally or globally distributed in aquatic environments (Fig. 4). For example, we report here a novel species, for which we propose the name “*Ca*. Aquidulcis frankliniae” (“Chloroflexi”), that is widely distributed geographically but restricted to freshwater environments. This species (and genus) is clearly distinct from its closest relative (“*Ca*. Limnocylindria sp”) based on phylogenetic reconstruction (Fig. S2-B) and AAI (71.85%). However, it would have remained cryptic if using 16S rRNA sequences alone, with a sequence identity of 98.4% between the two genera; a phenomenon previously observed for a few other bacterial taxa [32].

In order to evaluate preference (geographic or environmental), we devised a metric to compare expected and observed presence frequencies (Fig. 5) based on the observation that “presence” can be confidently assessed at the species level (95% ANI) and 0.05 p-value significance given a genome sequencing breadth of at least 10% [51]. All detected species appeared to have a preference for either freshwater or saltwater, or were too scarcely present to determine preference. Moreover, our method was able to distinguish species present in the estuaries that appeared to be adapted to freshwater, seawater, or displaying a preference specifically to estuaries (Figs. 4, 5). This clear differentiation likely reflects the large number of genomic adaptations required for a freshwater/seawater transition (*e.g.*, see [72, 73]).

Interestingly, we identified a statistically significant association between the ecologic range of gspp (in terms of habitat range and geographic distribution) and their genome size and coding density, indicating that more cosmopolitan gspp exhibit smaller, more compact genomes (Fig. S5). At first glance, this result might appear unexpected when considering that bacteria with more flexible and versatile metabolisms (multiple amenable carbon sources, detoxification mechanisms, or micronutrient scavenging capabilities) tend to have larger genomes, on average, and thus, are expected to colonize a higher number of ecological niches [74]. However, metabolic flexibility is also associated with fitness costs through the impact on growth rates, which may hinder the wider distribution across different habitats and long geographic distances. Indeed, we observed a phylogenetically-dependent negative association between estimated minimum generation time and ecologic range, indicating that (at short evolutionary distances) faster maximum growth facilitates more cosmopolitan distributions. These results support the hypothesis that benefits from metabolic flexibility provided by larger genomes could be superseded by the cost on fitness of replicating a longer genome and thus, longer generation times, on average [75, 76]. This idea has been formally described as the Black Queen Hypothesis (BQH), positing that genome reduction confers an inherent selective advantage to bacteria [77]. BQH has been used to explain the genome reduction of taxa with high population numbers (small effective population sizes are typically used to explain genome reductions such as those observed in endosymbionts), as observed in the marine members of the genera “*Ca*. Pelagibacter” and *Prochlorococcus* [77, 78] as well as in freshwater microorganisms including members of “Actinobacteria” and “Chloroflexi” [6, 7]. Here, we show that the effects predicted by BQH may be observed across *Bacteria*. Moreover, BQH implies the reliance of cosmopolitan bacteria on cheating: unilaterally using common goods such as secreted metabolites and extracellular proteins. In contrast, it has been previously proposed that cooperative pathogenic bacteria, not cheaters, have wider host ranges [70]. We found that, in our collection, there is a negative correlation between the fraction of extracellular proteins and all evaluated ecologic range metrics, further supporting BQH.

A consequence of BQH pervasiveness is that its effects should be observable in entire communities, not only in specific populations. While this prediction remains speculative, it is worth noting that selection for generalists, an increase in functional diversity, and faster growth rates have been observed in prokaryotic communities after a strong disturbance without an associated increase in average genome sizes [79]. However, note that these observations are based on genomes from samples geographically and environmentally restricted, and the generalization to other aquatic systems remains speculative.

Finally, most of the analyses described above required a reliable estimation of the relative abundance of genomospecies in each dataset. However, estimating MAG abundances in metagenomes is encumbered by: (1) genome incompleteness and imperfect estimations of completeness, (2) genome contamination and time-consuming and subjective contamination identification, and (3) microdiversity potentially confounding gene-content diversity with technical artifacts like non-overlapping assemblies. We applied a novel approach to estimate MAG abundance in metagenomes that sidesteps these limitations. Two key corrections include (1) truncation of sequencing depth before averaging to exclude highly conserved regions (overestimating depth), regions with gene-content micro-diversity (underestimating depth), and contamination (both); and (2) normalization of sequencing depth by genome equivalents in the metagenome, allowing relative abundance estimates. Note that this approach aims to estimate the **relative abundance of the species in the community** (*i.e.*, number of cells per total cells), not the more common metric of **relative abundance of sequenced DNA** which is affected by genome sizes [80]. Our abundance estimates correlated well with read counts normalized by metagenome size and genome length (RPKM [81]), while revealing an expected error of about 0.26 percent points in the simpler metric of read fraction (significantly correlated with completeness and N50, unlike our estimate) as well as about 1/3 of non-zero read fractions being potentially spurious. The metric introduced here has several advantages with respect to RPKM: (1) it is expressed in units of community fraction and, thus, can be readily interpreted as relative abundance, (2) it is robust to spurious high-depth regions due to highly conserved loci, and (3) the difference between zero and non-zero values is meaningful, as it corresponds to the tipping point for statistically significant presence.

In conclusion, we present methodological advances for the generation and study of MAGs derived from sets of related metagenomic datasets, and apply them to interconnected lakes and estuaries along the Chattahoochee River. This collection represents a valuable repository for the study of freshwater communities, and the methods introduced here are widely applicable to other metagenomic collections and environments. In addition, we show that cosmopolitan gspp tend to display smaller genomes with a phylogenetically-dependent association with faster growth rates, potentially reflecting the effects of the Black Queen Hypothesis.

### Data Availability

High-quality bins, distances, and other taxonomic analyses are available at http://microbial-genomes.org/projects/WB_binsHQ. Assembled genomes were also deposited in the NCBI GenBank database under BioProject PRJNA495371. All metagenomic datasets from the Chattahoochee samples are available in the NCBI SRA database as part of the BioProject PRJNA497294. Additional metadata on the provenance of sets in the iterative subtractive binning is also available as BioSamples SAMN10265471-SAMN10265528.

## Acknowledgments

We thank James R. Cole, James M. Tiedje (Michigan State University), William B. Whitman (University of Georgia, Athens), and Ramon Rosselló-Mora (Mediterranean Institute for Advanced Studies) for useful discussions on the methodology applied and ecologic implications of our results. We also thank Brittany Suttner, Yuanqi Wang, and Carlos A. Ruiz Perez (Georgia Institute of Technology) for support on fieldwork and sample processing, and Janet Hatt (Georgia Institute of Technology), and Patrick Chain (Los Alamos National Laboratory) for support on sample sequencing.

## Funding

This work was partly funded by the US National Science Foundation, awards #1241046 and #1759831 (to KTK).

## Competing Interests

The authors declare no competing interests.

## Supplementary Online Material

**Figure S1:** Total Community Diversity captured by the WB MAGs collection. **(A)** Shannon diversity (*H’*) estimated on the WB genomospecies abundance profiles (circles) and linear model of *H’* by *N_d_* (Nonpareil sequence diversity index; dashed line). The expected diversity based on *N_d_* is presented for reference (solid line) as derived previously [3]. Grey bands indicate the 95% confidence interval of both linear models. **(B)** Residuals of the observed *H’* with respect to the expected value; *i.e.*, distances between the circles and the solid line in panel A. **(C)** Total added abundance of the entire MAG set as a fraction of the community (y-axis) by *N_d_*. Note the significant positive linear correlation in panel B and the significant negative correlation in panel C, indicating that the more diverse communities (larger *N_d_*) have poorer diversity coverage by the WB MAG set (larger residuals in B, smaller total community fraction in C).

**Figure S2:** Phylogenetic reconstructions of two groups of MAGs and relatives. **(A)** Genome representatives from seven phyla within the “Terrabacteria” group, including “*Ca*. Elulota” proposed here. This species phylogeny represents a coalescent-based reconstruction on the trees of 82 genes [38–40]. **(B)** Representatives from four phyla within the “Terrabacteria” group, emphasizing the class “*Ca*. Limnocylindria” in the phylum “Chloroflexi”. This class includes two genera: “*Ca*. Limnocylindrus” (emended here) and “*Ca*. Aquidulcis” (proposed here). This species phylogeny obtained by coalescent-based reconstruction based on the trees of 67 genes. In both panels the genomes derived from metagenomes (MAGs) are prefixed with a label in squared brackets indicating the study from where they were derived, including: A13 [82], B15 [53], M18 [6], P18 [83], and Z18 [84], as well as the current study (This study) or publicly available data currently missing a published manuscript (Unpub). Genomes including a 16S rRNA gene sequence are marked with an asterisk.

**Figure S3:** Histograms of 80% central truncated average sequencing depth (TAD) all WB genomospecies in all Chattahoochee samples (top) and all other samples (bottom). Genomospecies with non-zero TAD (*i.e.*, a sequencing breadth > 10%) were considered confidently present if TAD was at least 0.01X (black), and uncertain otherwise (grey). The values of absent, uncertain, and present also correspond to the values in Fig. 4.

**Figure S4:** Phylogenetic diversity of the WB collection of MAGs in the context of best matches to other genomic collections **(A-B)** and the reference collection of genomes in PhyloPhlAn **(C-D)**. In panels A and C, the X-axis corresponds to the phylogenetic distance (branch lengths, bottom scale) at which the tree is cut into clades (numbers on top). These clades are then classified as containing only genomes from the WB collection (blue), only genomes from the reference database (grey), or both (blue and grey pattern). In panels B and D, the overall fraction of the tree (in branch lengths) covered by each category is summarized as Faith’s Phylogenetic Diversity (bar graph) in order to estimate the phylogenetic gain (right). Additionally, panel A includes an approximated phylogenetic calibration for the taxonomic ranks of species, genus, class, and phylum (vertical dashed lines). Each of these points also include the number of taxa only represented in the WB collection (novel) and the total number of taxa formed at the calibrated point.

**Figure S5:** Metrics of Ecologic Range and their correlation with genomic signatures. The y-axes (rows) indicate the genomic signatures evaluated, with the summary histograms in the rightmost panels. Conversely, the x-axes (columns) indicate the ecologic ranges, with summary histograms in the bottom panels. Both the correlation statistic (Pearson’s R or Spearman’s ρ)and the corresponding p-value are shown underneath each panel, with significant correlations (p-value < 0.01) highlighted in green (positive) and red (negative). The ecologic range metrics evaluated (left-to-right) are: biome count breadth (out of 13 biomes), aquatic habitat count breadth (out of 5 habitats), unweighted Levins’ breadths of biome and aquatic habitat (natural units), weighted Levins’ breadths of biome and aquatic habitat (natural units), geodesic range (thousands of km), and latitude range (degrees). The genomic features evaluated (from top-to-bottom) are: coding density (%), expected genome size (Mbp), G+C content (%), frequency of the J COG category (%), minimum generation time (h), and optimal growth temperature (°C).

**Table S1:** Metagenomic datasets from water bodies along the Chattahoochee River used here, including accession numbers, sample attributes, physicochemical parameters, and sequencing attributes.

**Table S2:** Metagenome-Assembled Genomes (MAGs) from the WB collection and general statistics.

**Table S3:** Metagenomic datasets from other studies used here to determine geographic and environmental breadth or preference. The column “Collection name” corresponds to the collections in Fig. 1. The columns “Sample accession” and “Run” correspond to the BioSample and Run accessions in the SRA/ENA databases, respectively. Additional metadata is provided as derived from MGnify or the original studies. The column “Reference” indicates the source study, corresponding to references [13, 52–65], or “Unpublished” corresponding to data publicly available in the SRA/ENA databases but currently not linked to publications.

**Text S1:** Additional details on materials and methods used in the present study.

**Text S2:** Additional details on population abundance and community diversity estimation.

**Text S3:** Additional methodology, results, and protologues for novel lineages described here: “*Ca*. Elulimicrobium humile” gen. nov. sp. nov. and “*Ca*. Aquidulcis frankliniae” gen. nov. sp. nov.

