## Supplementary figures and images for "Iterative Subtractive Binning of Freshwater Chronoseries Metagenomes Identifies of over Four Hundred Novel Species and their Ecologic Preferences"

### Figure S1

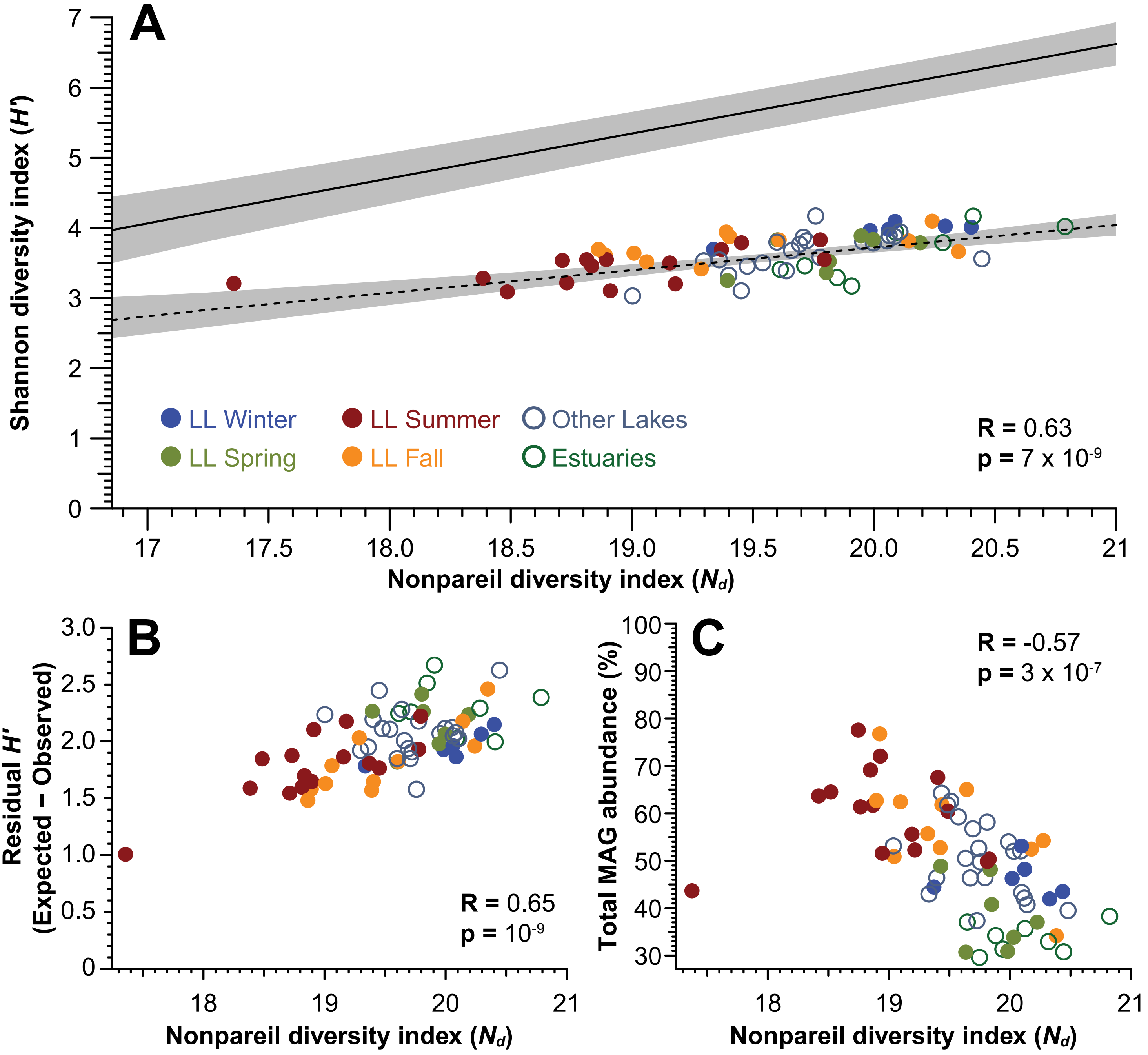

### Figure S2

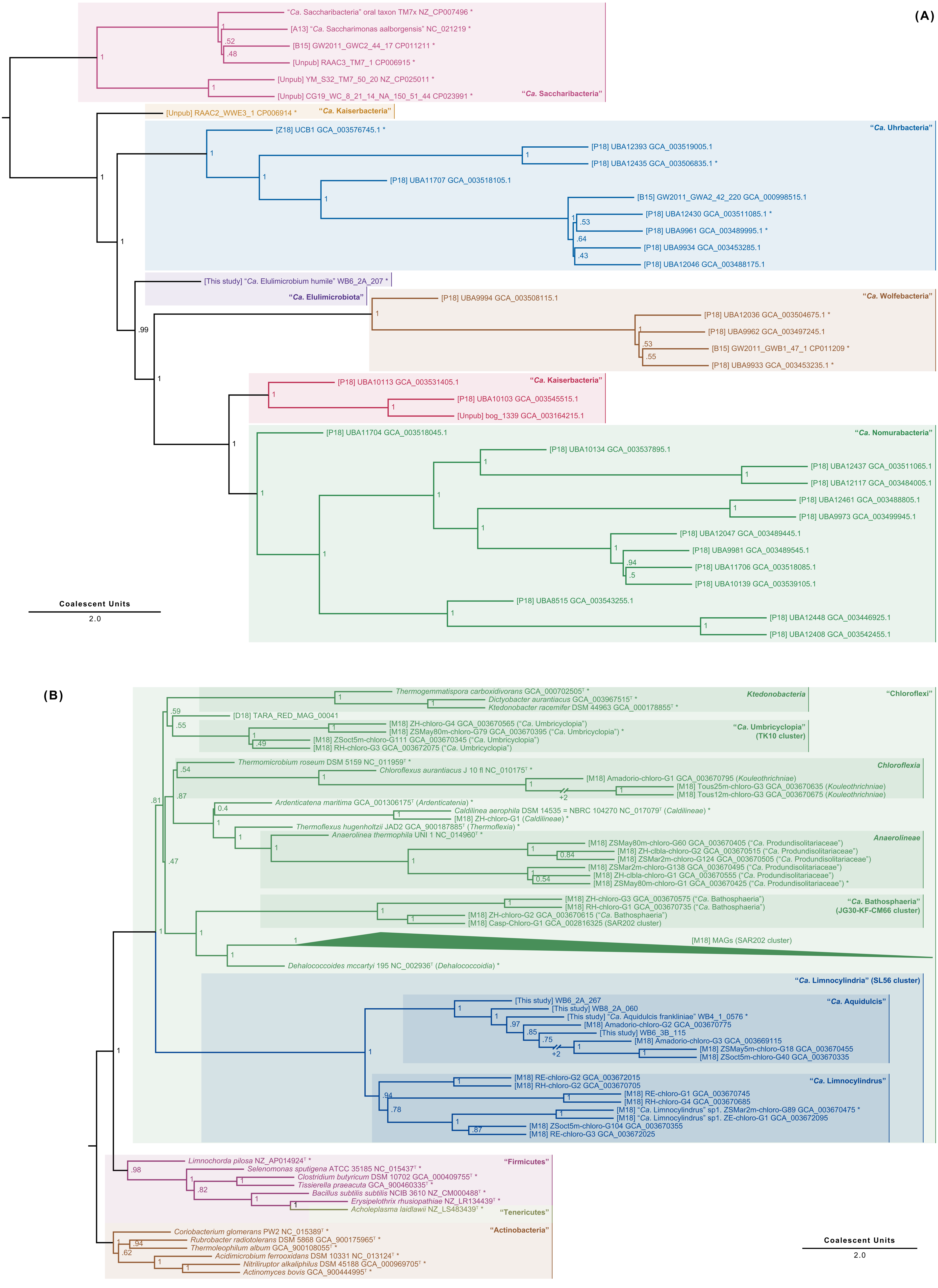

### Figure S3

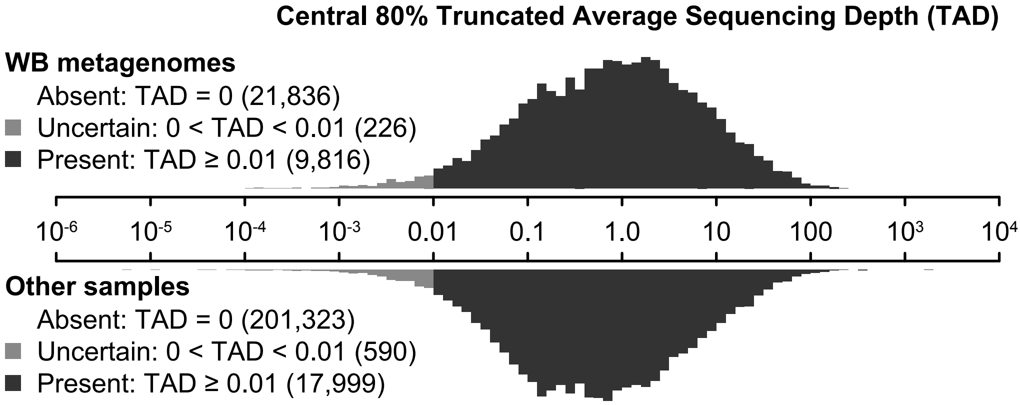

### Figure S4

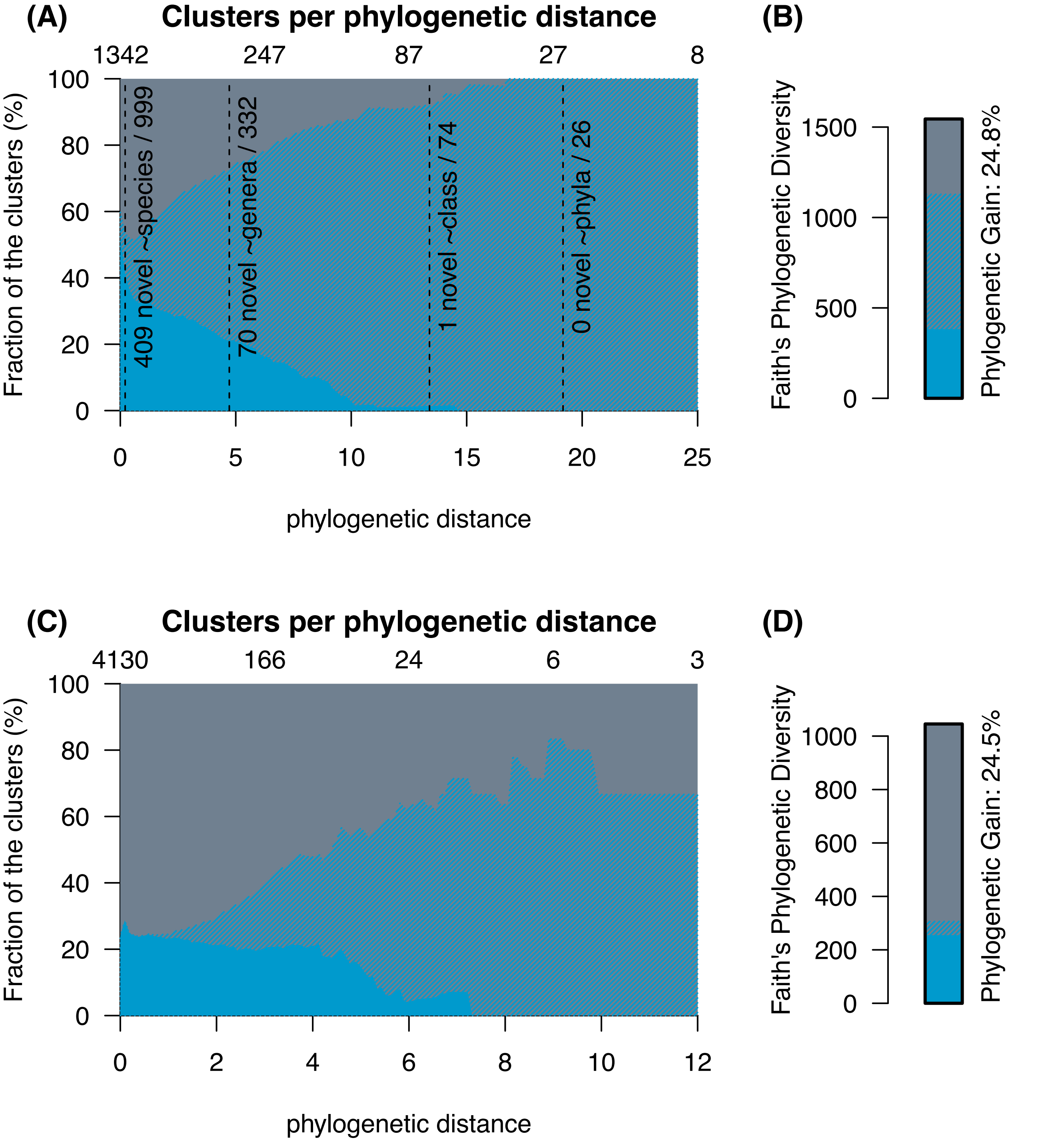

### Figure S5

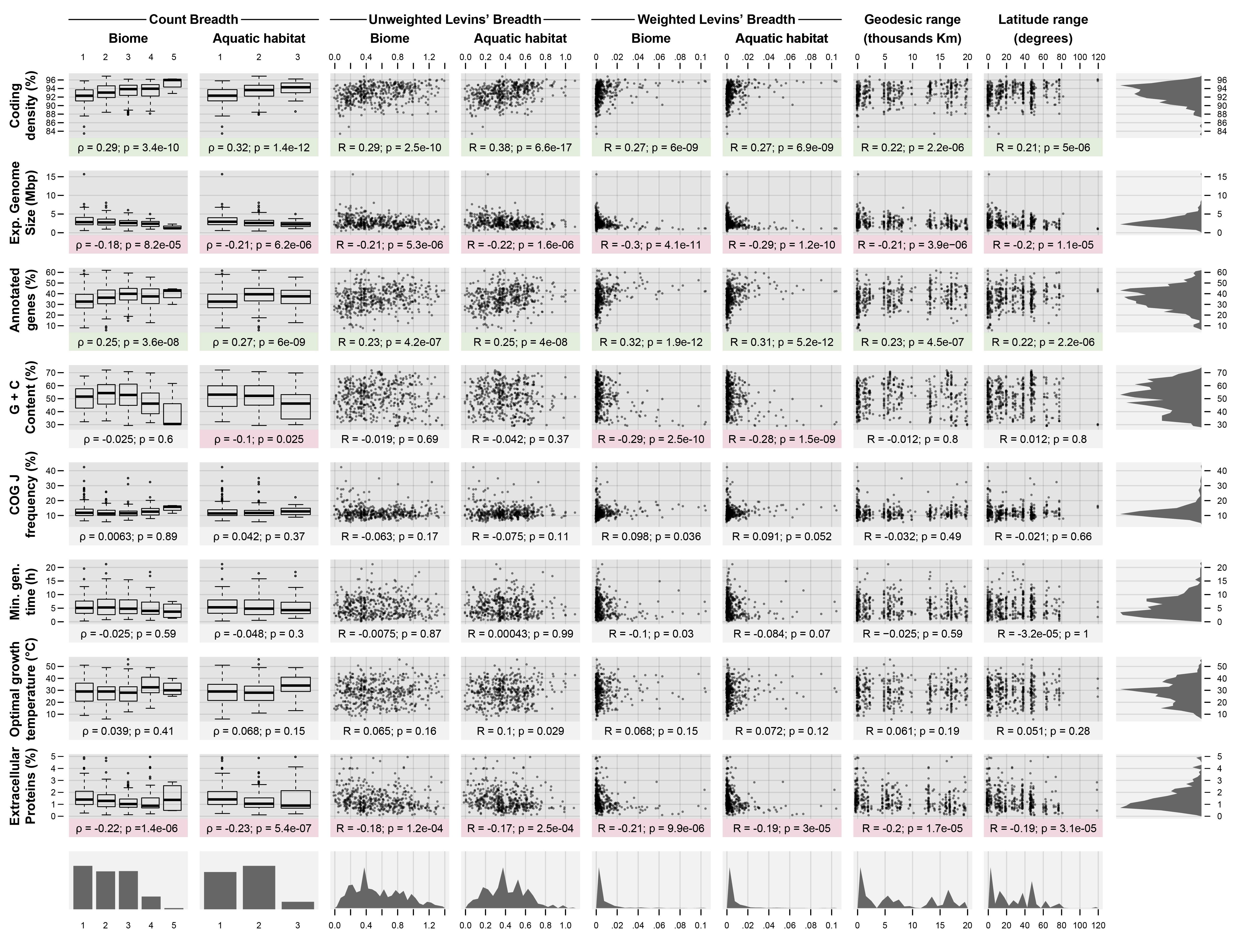
