## Supplementary material for "Iterative Subtractive Binning of Freshwater Chronoseries Metagenomes Identifies of over Four Hundred Novel Species and their Ecologic Preferences": Text S1

#### Supplementary Online Material S1: Extended methods

### Materials and Methods

##### Metagenomic Sequencing

DNA extraction was performed as previously described [16] with minor modifications. Briefly, frozen filters were fragmented and placed in microcentrifuge tubes with lysis buffer (50 mM Tris-HCl, 40 mM EDTA, and 0.75 M sucrose) and 1 mg/ml lysozyme, and incubated at 37°C for 30 min. Reactions were subsequently incubated with 1% SDS, 10 mg/ml proteinase K, and 150 μg/ml RNase for 4 h at 55°C in a rotating hybridization oven. DNA was extracted from the lysates with phenol and chloroform, precipitated with ethanol, and eluted in Tris-EDTA (TE) buffer. On average, DNA yield was 1.7 μg per liter of water filtered. One nanogram of DNA was used to prepare sequencing libraries using the Nextera XT Kit described by the manufacturer (Illumina). Libraries (9-11 pM) were sequenced using Illumina GA II, HiSeq, or MiSeq following manufacturer's protocols.

**Preference Score**

First, let us define $P_{GS}$, the binary matrix with elements $p_{ij} = \{1, 0\}$ indicating presence or absence of the gsp $i \in G$ (row) in sample $j \in S$ (column; see § Abundance and Alpha Diversity above for the definition of presence). Next, we define the expected frequency of observations as follows:

$$F_{GS} = \left(\frac{\overrightarrow{p_{G\bullet}} + 1}{\|S\| + 2}\right)^T \left(\frac{\overrightarrow{p_{\bullet S}} + 1}{\|G\| + 2}\right)$$

Where $X^T$ is transposed $X$; $\overrightarrow{p_{G\bullet}}$ is the sum of rows in $P_{GS}$, *i.e.*, the number of samples in which each gsp is present; $\overrightarrow{p_{\bullet S}}$ is the sum of columns in $P_{GS}$, *i.e.*, the number of gspp present in each sample; and $\|S\|$ and $\|G\|$ are the total number of samples and gspp considered, respectively. Note that the element-wise interpretation of the definition above for $f_{ij}$, is simply the probability of observation of gsp *i* across samples multiplied by the probability of observation of any gsp in sample *j*, with probabilities estimated by additive smoothing (add-one pseudocounts). We then determine the observed and expected biases in the frequency of gspp, $B_G$, in a given test set of samples, $t \subset S$, as follows:

$$B_G^{obs}(t) = \frac{2\overrightarrow{p_{Gt}} - \overrightarrow{p_{G\bullet}}}{\sum_{i,j} P_{ij}}$$

$$B_G^{exp}(t) = \frac{2\overrightarrow{f_{Gt}} - \overrightarrow{f_{G\bullet}}}{\sum_{i,j} F_{ij}}$$

Where $\overrightarrow{p_{Gt}}$ is the sum of rows for the columns of $P_{GS}$ in $t$, *i.e.*, the number of samples of $t$ in which each gsp is present; $\overrightarrow{f_{Gt}}$ is the sum of rows for the columns of $F_{GS}$ in $t$, and $\overrightarrow{f_{G\bullet}}$ is the sum of rows in $F_{GS}$. Note that $B_G(t) = -B_G(t^{\complement})$, where $t^{C}$ is the complement of *t*. Finally, we define the preference score of gsp *i* in *t* as:

$$s_i^p(t) = \frac{b_i^{obs}(t)}{\left|b_i^{exp}(t)\right|}$$

Where $|X|$ is the absolute value of $X$ and $b_i(t)$ is the *i*-th element of the vector $B_G(t)$. Note that the resulting score is symmetric, *i.e.*, $s_i^p(t) = -s_i^p(t^C)$. Genomospecies preference for sample set $t$ was considered significant when $s_i^p(t) > 1$, and gsp preference for $t^C$ was considered significant when $s_i^p(t) < -1$. No clear preference was established for gspp with $1 \geq s_i^p(t) \geq -1$.
