## Supplementary material for "Iterative Subtractive Binning of Freshwater Chronoseries Metagenomes Identifies of over Four Hundred Novel Species and their Ecologic Preferences": Text S2

### Supplementary Online Material S2: Population Abundance and Community Diversity Estimation

#### Population Abundance Estimation

In this study we use an estimator of abundance devised to reduce the impact of genome completeness from draft-level MAGs and control for other artifacts. Our metric is based on **TAD** values (80% truncated average sequencing depth) normalized by genome equivalents. However, we also computed the more traditional metrics of mapped metagenome **read fraction** and **RPKM** (reads per kilo-base pairs per million reads) [1] for all representative MAGs in the WB collection in all metagenomic datasets produced in this study. For each metagenome, we estimated the abundances of all MAGs with each of the three metrics, and compared these profiles by linear correlation (Pearson's R) and root-square of the mean squared error (RMSE, when compatible units). The estimated abundances correlated well with metagenome read fractions (mean Pearson's R: 0.89, IQR: 0.88-0.93), a metric that only controls for metagenome sizes, but the two estimates differed by about 0.26 percent points (mean RMSE;

IQR: 0.19-0.32%), which can be significant for rare members of the community. Moreover, read fractions were significantly correlated with estimated genome completeness (R=0.15, p-value=$10^{-3}$) and N50 (R=0.11, p-value=0.015), potentially obfuscating biologically meaningful correlations between abundance and genomic signatures. This artifact was not observed for our estimate of abundance (R=0.03 and 0.05, p-values=0.55 and 0.26, respectively). Moreover, spurious matches resulted in 97% of read fractions > 0, whereas our estimate (ensuring sequencing breadth ≥ 10%) only resulted in 31% non-zero values. The correlation of abundances was almost perfect with the RPKM metric (mean R: 0.997; IQR: 0.997-0.999), a metric that controls for the genome length in addition to the metagenome size, but the units of RPKM cannot be directly interpreted as relative abundances of the total community.

#### Community Diversity Estimation

In order to identify potential biases in the abundance profiles, we compared the expected alpha diversity in the community with the observed diversity in the MAGs abundance profile. Total community alpha diversity was estimated using the metagenomic read-based Nonpareil diversity index ($N_d$) and projected to the expected Shannon diversity index (*H’*) [2]. The diversity captured in the MAGs abundance profile was directly calculated as *H’* of the relative abundance vectors. In general, we observed a significant correlation between observed and expected alpha-diversity (Pearson’s R=0.63, p-value=7e-9; Fig. S1-A). However, we also observed larger residuals in more diverse communities, indicating that

MAGs capture a disproportionally larger fraction of less diverse communities (Pearson's R=0.65, p-value=1e-9; Fig. S1-B). This is also confirmed by the negative correlation between the total fraction of a community captured by MAGs and their Nonpareil diversity (Pearson's R=-0.57, p-value=3e-7; Fig. S1-C).
