## Supplementary material for "Iterative Subtractive Binning of Freshwater Chronoseries Metagenomes Identifies of over Four Hundred Novel Species and their Ecologic Preferences": Text S3

### Supplementary Online Material S3: Taxonomy of novel lineages

#### Genome Phylogeny

The phylogenetic reconstructions for specific taxa followed the same methodology as for the ASTRAL tree in the main text with one modification: all marker proteins present in at least 70% of the genomes were included, whereas the ASTRAL tree in the main text used marker proteins present in 80% of the genomes.

#### Novel Lineages

Using taxonomic information derived from the TypeMat, NCBI_Prok, UBA, and GCE collections, we classified monophyletic clades by phylum and, in the case of "Proteobacteria", by class (Fig. 3). The bacterial MAGs in this study span: "Proteobacteria" (194 gspp in 8 classes), "Dependiatiae" (4 gspp), "Acidobacteria" (3 gspp), "Spirochaetes" (1 gsp), the FCB Group (70 gspp in 4 phyla), the PVC group (72 gspp in 3 phyla), "Terrabacteria" (100 gspp in 7 phyla), and "Patescibacteria" (14 gspp in 6 phyla). From these, one phylum in

“Patescibacteria”, one phylum in “Terrabacteria”, and three classes in “Proteobacteria” are not named yet (novel).

In the “Patescibacteria” group, a novel phylum is uniquely represented by **WB6_2A_207** from our collection (76% completeness, 1.8% contamination, complete 16S rRNA gene detected with lowest RDP classification: Bacteria, 100% confidence). The gsp of WB6_2A_207 was detected in the August 2012 sampling of Lake Harding (LH_1208A) with an estimated relative abundance of 0.12%, and in 19 samples between July and October of all sampled years (2010-2015) from lakes Lanier, West Point, and Eufaula (*i.e.*, only detected upstream of Lake Eufaula, inclusive) with estimated abundances consistently below 0.06%. This gsp was not detected in any of the metagenomic datasets from other locations (Fig. 4). The fact that this gsp was detected in all six sampled years, in the four northernmost lakes but not in any of the downstream locations or any other geographic locations, and displays seasonal frequency (only late summer and early fall), indicates that this group is regionally endemic. We propose here that the species represented by WB6_2A_207 be named “*Candidatus* Elulimicrobium humile” gen. nov. sp. nov., after the month of harvest in the Akkadian calendar (Elūlu) typically corresponding to the annual period of detection for this group (August-September), and the Latin word *humile* (humble, low) indicating its consistently low abundance and restricted geographic range. This species establishes and serves as the type species for the taxonomic lineage up to “*Ca*. Elulimicrobia” cla. nov., in addition to “*Ca*. Elulimicrobiota” phy.

nov. within the group “Patescibacteria”, contingent upon the adoption of a formal phylum definition [1, 2]. A more detailed phylogenetic reconstruction based on coalescent reconciliation of 82 maximum likelihood gene trees including other representatives from the “Patescibacteria” places this novel group as sister clade to the group including “*Ca*. Wolfebacteria” (16S rRNA gene local identities against WB6_2A_207: 70.6-72.4%; AAI: 39.4-39.9%), “*Ca*. Kaiserbacteria” (no 16S rRNA gene sequences available; AAI: 38.3-38.8%), and “*Ca*. Nomurabacteria” (no 16S rRNA gene sequences available; AAI: 38.7-39.9%), although it might result in higher identities to (more distant) members of the phylum “*Ca*. Uhrbacteria” (16S rRNA gene identities: 69.6-74.5%; AAI: 38.6-39.3%) due to uneven branching patterns (Fig. S2-A).

In the “Terrabacteria” group, a distinct clade branching sister to “Chloroflexi” and close to “Firmicutes” was formed by four gspp in our collection, which could correspond to a novel order within “Chloroflexi” or a novel phylum. One genome from this set in our collection, **WB4_1_0576**, had a complete 16S rRNA gene sequence (lowest RDP classification: Bacteria, 100% confidence) and an estimated completeness of 83% and no detectable contamination. The closest relative detected was the MAG **RED_MAG_00041** (AAI against WB genomes in this clade: 40.0-40.5%), derived from Tara Ocean samples from the Red Sea [3], and had no detected 16S rRNA gene (not assembled or binned), 78% completeness, and 0.9% contamination. The gsp of WB4_1_0576 was detected in 62% of all metagenomic samples in this study, representing all sampling sites except the East Point estuary, although it was detected in the neighboring Apalachicola estuarine location in three non-consecutive years (2010, 2013, and 2015). The maximum estimated relative abundance only surpassed 0.1% of the community in Lake Seminole (0.3%), followed by the adjacent locations of Lake Eufaula (North, 0.033%) and the Apalachicola estuary (South, 0.031%). No clear seasonal pattern was observed for this gsp in Lake Lanier, and seasonal resolution was not available for Lake Seminole. Additionally, this gsp appears to be globally distributed, but with estimated relative abundances only as high as 0.2% in Lake Mendota (Madison, Wisconsin, USA; BioSample SAMN05422235) [4], 0.1% in Lake Ontario (Ontario, Canada; BioSample SAMN05421521), 0.05% in the Xiangxi River (Hubei province, China; BioSample SAMEA4051564) [5], and 0.04% in Lake Erken (Sweden; BioSample SAMEA2241989) [6]. It was also identified in the Amazon River (Brazil; BioSample SAMN02924045) although at very low sequencing depth (0.0013X TAD) corresponding to only 0.001% of the community. Therefore, this group was only confidently identified in lake samples and in the Apalachicola estuary, revealing a strictly freshwater lifestyle and relatively low *in-situ* abundances. Further inspection into recently published freshwater MAGs revealed that close relatives to the group represented by this genome have been previously identified in other lakes around the world, and likely belong to the class "*Ca*. Limnocylidria" (also referred to as SL56 cluster) from the phylum "Chloroflexi" [7] (Fig. S2-B). Importantly, the proposed name for this class ("*Ca*. Limnocylindria") did not designate a type species for the genus (lowest proposed rank) nor did it specify type material. Therefore, we propose emending the description by establishing the genome ZSMar2m-chloro-G89 (only one in the clade from the original publication with complete 16S rRNA gene sequence) as type material. In addition, we propose that the species represented by WB4_1_0576 be named “*Candidatus* Aquidulcis frankliniae” gen. nov. sp. nov., after the Latin phrase *Aqua dulcis* (freshwater) and the late chemist Rosalind Elsie Franklin, whose work was pivotal on understanding the structure of DNA. All other genomes in our study (WB collection) within the same phylum belong to the same genus on the basis of AAI. The closest relatives, WB8_2A_060 (AAI: 90.9%) and WB6_3B_115 (AAI: 90.0%), displayed similar biogeographic patterns to WB4_1_0576. In contrast, the more distant relative within the genus (WB6_2A_267, AAI: 84.1%) had a more restricted distribution, detected only in eight samples from the Chattahoochee River, and in only two samples of the additional surveyed sites with abundances below 0.001% in Lake Ontario (BioSample SRR3990180) and Lake Mendota (BioSample SAMN05422235). It displayed intermediate abundances in Lake Eufaula, where it was detected in September 2014 (estimated relative abundance: 0.1%) and October 2015 (replicate estimated abundances: 0.5%, 0.5%, and 0.6%), and abundances below 0.1% in the Apalachicola estuary and lakes Harding and West Point. The genomes in the genus “*Ca*. Aquidulcis” proposed here displayed AAI values against the “*Ca*. Limnocylindria” genome ZSMar2m-chloro-G89 between 70.7 and 71.9% (AAI against WB4_1_0576: 71.85%). Interestingly, the 16S rRNA genes of WB4_1_0576 and ZSMar2m-chloro-G89 displayed high sequence identity (98.4%), despite their relatively low AAI.

In the phylum “Proteobacteria”, we detected members of three yet-unnamed classes. The first one, a clade sister to *Gammaproteobacteria* (around 2 o’clock in Fig. 3), was formed by four gspp in our collection (WB7_3xA_002, WB4_4_611, WB7_3xI_023, and WB7_3xI_002), two Tara MAGs (MED_MAG_00110 and ION_MAG_00050), and two UBA MAGs (UBA4577 and UBA3005). None of the above had detectable 16S rRNA gene fragments. The second one, a clade branching between the class *Betaproteobacteria* and the order *Acidiferrobacterales*, was formed by two gspp in our collection (WB6_4_032 and WB5_5_101), two UBA genomes (UBA868 and UBA2133), and a Tara MAG (PSW_MAG_00045). From these, only one genome had 16S rRNA gene fragments detected (WB6_4_032, 4 partial 16S sequences), but these were likely the result of contamination as the classification of the fragments spanned different bacterial phyla. According to the reconstructed phylogeny, it is likely that this clade forms a single class together with the order *Acidiferrobacterales*, represented by its type species *Sulfurifustis variabilis*. However, the latter was originally classified as part of *Gammaproteobacteria* [8], a classification that is not congruent with our phylogenetic reconstruction. Instead, we propose this group be affiliated to a separate class including the genera *Acidiferrobacter* and *Sulfurifustis* [8, 9], as well as any other genera represented by the MAGs listed above. Finally, a monophyletic group sister to the clade including *Gammaproteobacteria*, *Betaproteobacteria*, and both yet-unnamed classes described above, was formed by three gspp in our collection

(WB4_3_0799, WB4_2_1502, and LLD_282). None of these had detectable 16S rRNA gene fragments, but one of these genomes was of high quality (LLD_282: 84% completeness, no contamination detected).

**Description of "*Candidatus* Elulimicrobium" gen. nov**

*Elulimicrobium*, [E.lu.li.mi.cro'bi.um. N.L. neut. n. *elulu* from Akkadian n. *elūlu*, and from there to Hebrew n. Elul (אלול), the month of harvest in the Akkadian calendar typically corresponding to the annual period of detection for this group (August-September); N.L. neut. n. *microbium*, microbe]. The genus is established on the basis of phylogenetic reconstruction including the type species "*Ca*. Elulimicrobium humile" and other members of the group "Patescibacteria" (Fig. S2-A). The type species is "*Ca*. Elulimicrobium humile".

**Description of "*Candidatus* Elulimicrobium humile" sp. nov**

*E. humile*, [hu.mi'le. L. neut. adj. *humile*, humble, low]. This species displays a consistently low abundance when detected between the months of July and October, and a geographic range restricted to the tree northernmost lakes on the Chattahoochee River (USA): lakes Lanier, West Point, and Eufaula. According to the genome assembly, WB6_2A_207 is 3.79 Mbp long with 1,298 contigs (N50: 4 kbp) and 34.3% G+C content. It includes 4,108 predicted proteins as well as 90 non-coding RNA loci (87 tRNA, 1 rRNA, and 2 tmRNA). The estimated coding density is 94.1%. Based on genome annotation, this organism is predicted to have sucrose and oxidase activity and the capability of gelatin and arginine dihydrolase activity. The type material is the genome WB6_2A_207, deposited in the NCBI database with Genome Accession RGCK00000000 and BioSample SAMN10223143.

**Description of "*Candidatus* Elulimicrobiaceae" fam. nov.**

*Elulimicrobiaceae* [E.lu.li.mi.cro.bi.a.ce'ae. N.L. neut. n. *Elulimicrobium* referring to the type genus of the family "*Ca*. Elulimicrobium"; *-aceae* ending to denote a family; N.L. fem. pl. n. *Elulimicrobiaceae* the "*Ca*. Elulimicrobium" family]. The properties of the family are the same as for the representative genus "*Ca*. Elulimicrobium", which is the designated type genus of the family.

**Description of "*Candidatus* Elulimicrobiales" ord. nov.**

*Elulimicrobiales* [E.lu.li.mi.cro.bi.a'les. N.L. neut. n. *Elulimicrobium* referring to the type genus of the order "*Ca*. Elulimicrobium"; *-ales* ending to denote an order; N.L. fem. pl. n. *Elulimicrobiales* the "*Ca*. Elulimicrobium" order]. The properties of the order Elulimicrobiales are the same as for the representative genus "*Ca*. Elulu", which is the designated type genus of the order.

**Description of "*Candidatus* Elulimicrobia" classis nov.**

*Elulimicrobia* [E.lu.li.mi.cro'bi.a. N.L. fem. pl. n. *Elulimicrobiales* referring to the type order of the class "*Ca*. Elulimicrobiales"; *-ia* ending to denote a class; N.L. neut. pl. n. *Elulimicrobia* the "*Ca*. Elulimicrobiales" class]. The class is defined on the basis of comparative genome analysis on the genus "Ca. Elulimicrobium" with respect to currently determined genomes from the closest relatives within the “Patescibacteria” clade. “*Ca*. Elulimicrobiales” is the designated type order of the class.

**Description of “Candidatus Elulimicrobiota” phy. nov.**

*Elulimicrobiota* [E.lu.li.mi.cro.bi.o’ta. N.L. neut. pl. n. *Elulimicrobia* referring to the type class “*Ca*. Elulimicrobia”; -*ota* ending to denote a phylum; N.L. neut. pl. n. *Elulimicrobiota* the “*Ca*. Elulimicrobia” phylum]. The phylum is defined on the basis of comparative genome analysis on the genus “Ca. Elulimicrobium” with respect to currently determined genomes from the closest relatives within the “Patescibacteria” clade, contingent upon the adoption of the rank of phylum [1, 2]. “*Ca*. Elulimicrobia” is the designated type class of the phylum.

**Emendation of “*Candidatus* Limnocylindrus”**

The properties of the species are as given by Mehrshad *et al*, 2018 [7], and the designated type material is ZSMar2m-chloro-G89, deposited in the NCBI databases with Genome Accession QWRJ00000000 and BioSample SAMN09724413.

**Description of “*Candidatus* Aquidulcis” gen. nov**

*Aquidulcis*, [A.qui.dul’cis. L. fem. n. *aqua*, water; L. fem. adj. *dulcis*, sweet; N.L. fem. n. (N.L. fem. adj. used as a substantive) *Aquidulcis*, an organism of freshwater]. The genus is established on the basis of Average Amino Acid

Identity (AAI) and phylogenetic reconstruction using comparative genomics of the genus representatives and representatives of "*Ca*. Limnocylindrus" (Fig. S2-B). Heterotrophic aerobic bacteria with genomic capability for reductive dehalogenation but not sulfite oxidation. The type species is "*Ca*. Aquidulcis frankliniae".

**Description of "*Candidatus* Aquidulcis frankliniae" sp. nov**

*A. frankliniae*, [frank.li'ni.ae. N.L. fem. gen. n. *frankliniae*, of Franklin, in honor of the English chemist Rosalind Elsie Franklin, whose work was pivotal on the understanding of the DNA structure]. Aerobic photoheterotrophs encoding rhodopsins tuned for green-light absorption. The genome WB4_1_0576 has 1.2 Mbp in length in 154 contigs (N50: 15.5 kbp) with 61.9% G+C content, including 1,272 predicted proteins, as well as 45 tRNA and 3 rRNA loci. The estimated genome coding density is 95.5% and the complete genome is estimated to be 1.5 Mbp. Based on genome annotation, it is predicted to have bacillus or coccobacillus morphology and possibly stain Gram positive. The type material is the genome WB4_1_0576, deposited in NCBI databases with Genome Accession RFPZ00000000 and BioSample SAMN10222820.
